# Alpha oscillations and aperiodic neural dynamics jointly predict visual temporal resolution, confidence, and dependence on prior experience

**DOI:** 10.64898/2026.01.05.697694

**Authors:** Gianluca Marsicano, Michele Deodato, David Melcher

## Abstract

Perception requires integrating sensory input over time to construct coherent experiences. Alpha oscillations have been proposed to define the temporal resolution of perception, yet empirical evidence remains inconsistent. Here we combined a sustained visual integration paradigm with resting-state EEG to investigate how oscillatory and aperiodic neural dynamics jointly shape temporal perception. Participants (n=83) viewed alternating gratings that varied in alternation speed, producing the perception of either a fused plaid (integration) or two interchanging gratings (segregation). Faster individual alpha rhythms were associated with narrower temporal integration windows, and a steeper aperiodic spectrum predicted greater perceptual precision. Moreover, slower alpha frequencies and flatter neural spectra were associated with stronger reliance on prior judgments, suggesting reduced sensory precision and increased weighting of recent experience. Individuals with faster alpha rhythms showed a higher general confidence tendency, but a reduced influence of trial-by-trial changes in temporal evidence on their confidence ratings. Together, these findings show that perceptual interpretation, confidence, and the influence of previous experience on temporal integration can be jointly predicted by alpha rhythms and aperiodic neural dynamics. Mechanistically, faster alpha cycles and lower neural variability may enhance perceptual resolution by generating more precise sampling frames per time unit, leading to finer temporal perception, reduced reliance on prior experience, and greater confidence.

## Introduction

Temporal integration is a fundamental operation in vision: it allows the nervous system to accumulate sensory evidence over brief time intervals so that perception remains robust despite internal noise and external uncertainty^1^. Yet, we still lack a clear account of how the brain sets the temporal window within which sensory signals are bound. In other words, how does the system decide whether two sequential inputs should be integrated or segregated?

A prominent proposal is that rhythmic activity in the alpha band (∼8–13 Hz) provides a temporal scaffold for perception. On this view, alpha oscillations define temporal excitation windows that allow for the integration and accurate processing of sensory information^2–5^. Accordingly, faster individual alpha frequencies (IAFs) would enable finer temporal resolution, whereas slower alpha rhythms would enlarge the integration window and promote binding. However, despite its conceptual appeal, the relationship between alpha oscillations and perception remains debated. Empirical evidence has been mixed, with several recent studies reporting null or even contradictory findings, calling into question the generality of this framework^6,7^.

Several factors may underlie these inconsistencies, both methodological and theoretical. One methodological issue is that most previous studies have relied on variants of the *two-flash fusion task*, in which participants judge whether two brief, identical flashes presented in close succession are perceived as one or two distinct events^2^. Within the framework of rhythmic perception, it is hypothesized that stimuli falling within the same alpha cycle are bound into a single percept. However, this interpretation of two flash perception is not unambiguous: because each flash can occur at different phases of the ongoing alpha rhythm, one stimulus might simply be processed suboptimally, yielding an apparent fused percept that reflects detection failure rather than genuine temporal integration^2,8–10^. One way to address this challenge is to use paradigms in which the stimuli are shown on screen for several seconds^8,11^.

On a more theoretical level, there is disagreement over the fundamental mechanisms that link alpha oscillations to visual temporal perception. From a Bayesian perspective, for example, alpha frequency may shape perceptual inference by altering perceptual sensitivity or by adjusting top-down priors^6,12^. However, this framework often overlooks that such priors may be dynamically shaped by recent perceptual experience, which continuously updates internal models and biases the interpretation of incoming sensory information (i.e., serial dependence^13,14^). Critically, recent evidence shows that temporal binding mechanisms are not fixed within individuals but adaptively expand or contract based on prior perception, with events following a binding percept more likely to be experienced as a single unified event^15–17^. Interestingly, it has been hypothesized that serial dependence itself may be linked to alpha oscillations. One possibility is that these oscillations facilitate communication, binding current sensory input with recent perceptual history^14,18,19^. It is also plausible that the speed of alpha oscillations affects how strongly past experiences shape current perception. When IAF is slower, temporal resolution and sensory precision may decrease^18,19^, making the brain’s estimate of current sensory input more uncertain and thereby increasing reliance on prior experience to stabilize perception.

Another critical aspect that has often been overlooked in studies of temporal integration is its metacognitive dimension. Subjective confidence systematically influences pre- and post-perceptual processing across sensory domains^20–22^. Notably, decision confidence reflects an observer’s subjective certainty about a perceptual judgment and is generally shaped by the strength of the underlying sensory evidence, while remaining partly dissociable from objective perceptual performance^23–26^. Thus, confidence in temporal binding may reflect both the strength of current temporal evidence and the recent perceptual context in which that evidence is interpreted. Recent evidence further suggests that faster alpha rhythms may enhance perceptual resolution by generating more sampling frames per time unit, and that more frequent sampling of sensory input may yield a more faithful representation of external events^18,19^, thus enhancing perceptual and metacognitive abilities. Individual differences in alpha rhythms may therefore influence not only temporal resolution, but also the subjective confidence assigned to perceptual judgments. In sum, the relationship between alpha oscillations and temporal perception may be more multi-layered, resisting a simple, single mechanistic explanation.

Another critical consideration is that the focus on oscillations may ignore a link to aperiodic activity, since the complex nature of visual temporal perception likely depends on multiple neural mechanisms, not only rhythmic oscillations^5,27–29^. Neural activity exhibits a prominent aperiodic *1/f* component reflecting nonrhythmic brain dynamics^29–31^, and while there is growing evidence for the importance of taking aperiodic spectra into account^29,31,32^, exactly how and why it influences temporal perception is still being explored^5,27,29,33,34^. The aperiodic exponent, operationalized as the slope of the spectral power decay across frequencies, quantifies the broadband organization of neural activity^29,32^. It varies systematically with age^29,35,36^ and across neural states^29,37,38^, predicts individual differences in behavior^27,33,39–41^, and is altered across a broad range of clinical conditions^29,33,36,42–45^. Flatter spectra have been associated with greater excitation and irregular activity, whereas steeper neural spectra may reflect stronger inhibitory constraints and greater temporal organization^29,31,46–49^. The aperiodic cortical state may therefore shape sensory processing and how reliably rhythmic oscillations organize information over time^27,29,33,34^. Oscillations may determine when sensory information is sampled, whereas the broadband background may influence encoding reliability through neuronal gain and processing variability. Thus, lower neural variability may support stronger rhythmic coordination, whereas increased cortical excitation may weaken oscillatory temporal sampling^27,34,50^. Within this framework, recent findings indicate that the association between alpha rhythms and temporal integration becomes apparent only when aperiodic activity is modeled alongside oscillatory dynamics, with flatter neural spectra linked to increased variability in visual temporal processing and longer temporal integration windows^5,27,34^. Hence, oscillatory and aperiodic neural dynamics may jointly shape visual temporal integration, providing a more complete account of how the brain samples sensory input over time and potentially explaining recent null findings.

In this study, we investigated resting-state neural dynamics to characterize trait-level neural predictors of the multidimensional aspects underlying visual temporal perception, focusing on the potentially complementary contributions of resting-state alpha frequency and aperiodic spectral dynamics. Both resting-state measures have been consistently linked to visual temporal perception^2,3,27,40,45^ and, more broadly, to individual differences in cognition and perception^29,39,41,51^, while also showing substantial within-individual stability^27,29,41,52–57^. We therefore asked whether these trait-like neural characteristics were associated with individual differences across multiple aspects of visual temporal performance, providing a multilayered neural account of why temporal perception varies across individuals.

To address this challenge, we developed a novel variation of a sustained visual stimulus paradigm^8,33^, based on the perceptual fusion of consecutive visual patterns presented in a temporally extended stream (Fig. 1A). A large sample of participants (n = 83) viewed two low– spatial-frequency Gabor patches oriented in opposite directions (±45°) that alternated in the same spatial location in counterphase over one second. Rather than manipulating the interval between discrete flashes, we varied the duration of each Gabor presentation. This created a continuous visual flow in which shorter durations elicited the illusory perception of a single fused plaid (integration), whereas longer durations produced the perception of two alternating gratings (segregation). This design allowed us to estimate individual temporal integration thresholds while avoiding the confounds of traditional paradigms (e.g., detection failures), providing a more direct measure of perceptual integration^8,33^. An advantage of continuously presented stimulus streams that integration performance is not confounded with missing a rapidly presented stimulus. Before the behavioral task, we recorded resting-state EEG in both eyes-open and eyes-closed conditions to examine how oscillatory and aperiodic neural dynamics jointly shape visual temporal integration. Critically, we considered the influence of prior perceptual judgment and subjective confidence, relating each of these factors to periodic and aperiodic activity. Including all of these measures of temporal integration performance allowed us to test a mechanistic framework in which faster resting-state alpha oscillations within a steeper neural background predicted an enhanced rate and fidelity of visual sampling^5,18,19,27,34,58^, shaping how the visual system integrates sensory information over time.

**Figure 1.**
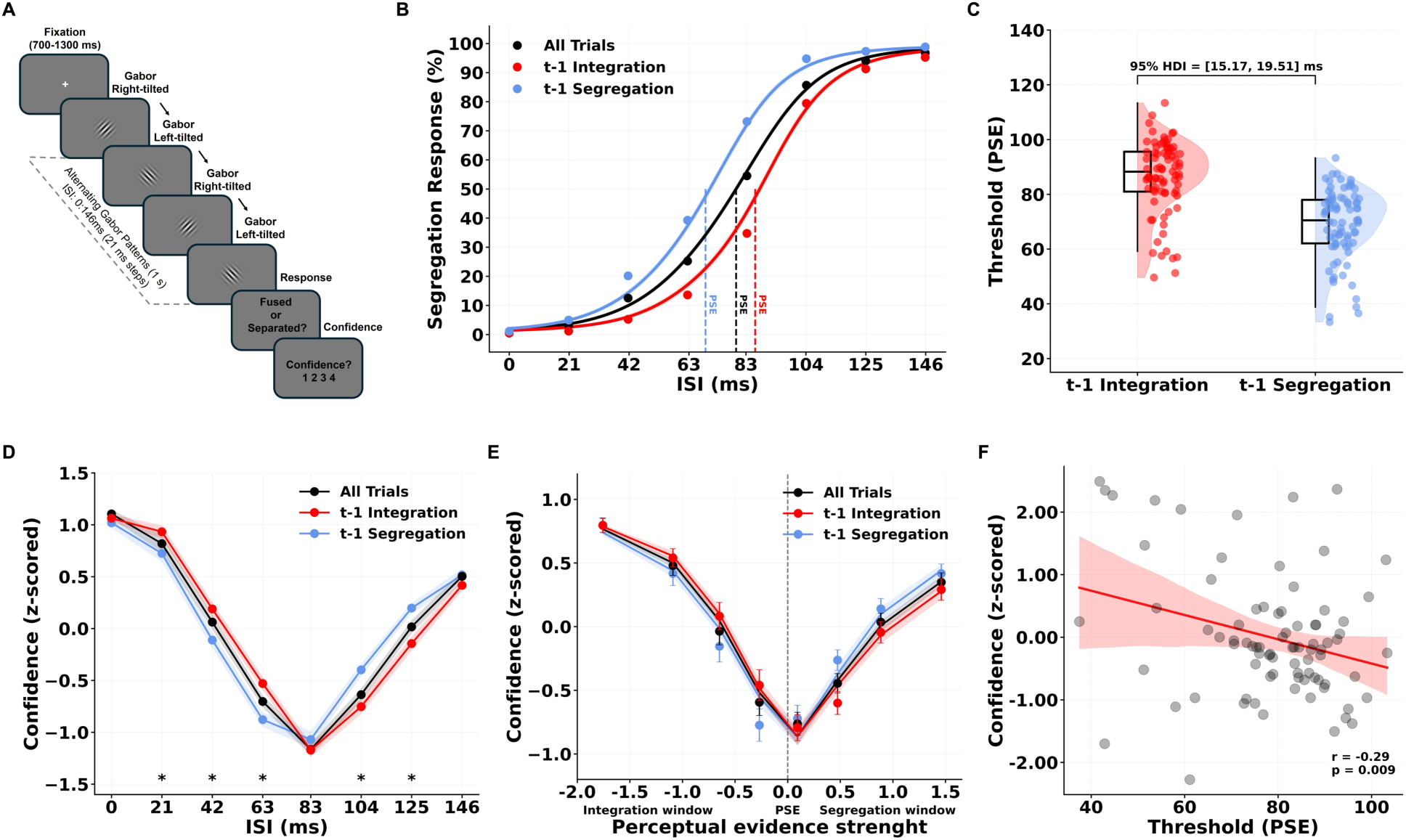
Visual temporal binding is dynamically shaped by sensory input, perceptual history and confidence. (A) Schematic depiction of the sustained-stream temporal integration task. On each trial, two low–spatial-frequency Gabor patches of opposite orientation (±45°) alternated in counterphase for ∼1 s with variable inter-stimulus intervals (ISIs, 0–146 ms, steps of 21 ms). Participants reported whether they perceived a fused plaid (integration) or separate gratings (segregation), followed by a confidence rating (1–4). (B) Psychometric functions showing the proportion of segregation responses as a function of ISI for all trials (black), trials following t–1 integration responses (red), and t– 1 segregation responses (blue). Shaded areas denote SEM. The Point of Subjective Equality (PSE) was higher following t–1 integration trials, indicating a broader temporal binding window. (C) Individual PSE values as a function of previous perceptual outcome. Integration on the preceding trial biased perception toward integration on the current trial, indicating a stronger influence of recent perceptual choice-history on temporal binding processes (serial dependence effect). Black boxplots indicate the median and interquartile range (IQR), and individual dots represent participant-level data. The bracket indicates the 95% high density intervals (HDIs) of the group level difference. (D) Mean z-scored confidence ratings as a function of ISI and t–1 judgment. Confidence decreased near intermediate ISIs and increased when temporal evidence clearly supported either integration or segregation. Confidence was higher after t–1 integration responses at short ISIs (21–63 ms) and higher after t–1 segregation trials at long ISIs (104–125 ms; q < 0.05, FDR-corrected), showing an interaction between prior experience and temporal evidence. (E) Mean z-scored confidence as a function of individualized perceptual evidence strength, quantified relative to each participant’s PSE, for all trials (black), trials preceded by integration (red), and trials preceded by segregation responses (blue). Confidence was lowest near the perceptual threshold (PSE = 0) and increased as evidence more strongly favored either integration or segregation. Confidence was additionally enhanced when current evidence was congruent with the perceptual outcome of the preceding trial, indicating that subjective confidence was jointly shaped by current perceptual evidence and recent perceptual history. (F) Across participants, mean confidence negatively correlated with temporal integration threshold (PSE; *r* = –0.29, *p* = 0.009), indicating that individuals with narrower integration windows (higher temporal acuity) reported greater confidence. Shaded regions in line plots and scatterplots represent 95% confidence intervals. Together, these results show that visual temporal binding and subjective confidence are dynamically shaped by recent perceptual experience, reflecting an adaptive interplay between sensory evidence and perceptual history.

## Results

### Visual temporal binding is dynamically shaped by recent perceptual experience

The sustained stream of visual patterns in our temporal integration task elicited the illusory perception of a single fused plaid (integration) at shorter inter-stimulus intervals (ISIs), whereas longer ISIs induced the perception of two alternating gratings (segregation; Fig. 1A, 1B). This continuous task replicates classical findings from two-flash paradigms while avoiding their confounds^8,33^, providing a more direct measure of perceptual binding. To characterize behavioural performance and its latent psychometric parameters, we modelled segregation responses across ISIs using hierarchical Bayesian psychometric modelling (see Methods). Recent evidence indicates that visual temporal processing is not fixed but is dynamically modulated by recent perceptual choice history (i.e., serial dependence^15–17^). Accordingly, the modelling also accounted for the influence of the perceptual outcome of the preceding trial: (i) t−1 integration, corresponding to a fused response on the previous trial, and (ii) t−1 segregation, corresponding to a segregation response on the previous trial. The model that best characterized behavioural performance included participant-specific effects of the preceding percept on both the psychometric threshold and slope, together with a symmetric lapse parameter accounting for stimulus-independent responses. The model therefore yielded three principal group- and participant-level components: the temporal threshold, or Point of Subjective Equality (PSE), the psychometric slope, and the lapse rate. The PSE reflects the temporal boundary between perceptual integration and segregation. The psychometric slope quantifies how abruptly and consistently perceptual reports transition from integration to segregation, reflecting both sensory and decisional sources of trial-by-trial variability^59,60^. Finally, the lapse parameter estimates rare stimulus-independent response errors (e.g., attentional lapses, motor errors), allowing these responses to be distinguished from variability in the PSE and psychometric slope.

The group-level temporal integration threshold was centered around ∼80 ms (PSE = 77.53 ms; 95% HDIs [74.58, 80.48]), closely aligning with the frequency of alpha-band oscillations. As expected, visual temporal processing was dynamically modulated by perceptual choice history. Participants were more likely to repeat the perceptual interpretation reported on the preceding trial: integration expanded the temporal binding window, whereas segregation contracted it. Following an integrated percept, the group-level PSE was 86.25 ms (95% HDIs [83.05, 89.33]), whereas following a segregated percept it was 68.82 ms (95% HDIs [65.58, 71.89]), resulting in a history-dependent PSE difference of 17.43 ms (95% HDIs_difference_ [15.17, 19.51]; Pr(ΔPSE > 0) > .999; Fig. 1B,C). In contrast, there was no evidence for a consistent effect of perceptual history on psychometric slope (95% HDIs_difference_ log-slope [−0.088, 0.104]), indicating that recent perceptual history shifted the temporal integration threshold without consistently altering the precision of the perceptual transition. Finally, the estimated lapse parameter indicated a low rate of stimulus-independent responses, with a group mean of 1.70% (95% HDIs [1.06%, 2.34%]). This indicates that stimulus-independent responses (e.g., attentional fluctuations or random responses) were rare and were unlikely to substantially bias the estimated PSE or psychometric slope.

In line with recent evidence^16,17^, these findings suggests that temporal integration mechanisms are flexible within individuals, adaptively expanding or contracting based on prior perceptual outcomes, reflecting a dynamic interplay between sensory evidence and perceptual history.

### Confidence dynamically adjusts with temporal evidence and prior perceptual experience

After each temporal binding judgment, participants rated their subjective confidence on a four-point scale. As expected, confidence followed the temporal clarity of sensory evidence: participants reported lower confidence at intermediate ISIs, corresponding to perceptually ambiguous trials near the temporal integration threshold, and higher confidence at shorter and longer ISIs, where temporal evidence more clearly supported integration or segregation decisions. Across participants, mean subjective confidence was inversely related to the temporal integration threshold (PSE), revealing a negative correlation between confidence and PSE (*r* = –0.29, *p* = 0.009; Fig. 1F). Thus, individuals with narrower integration windows (i.e., higher temporal acuity) reported higher overall confidence in their perceptual judgments. In contrast, mean confidence was not significantly correlated with the slope of the psychometric curve (*r* = 0.152, *p* = 0.18). We next examined whether confidence was influenced by recent perceptual history. Trial-wise confidence ratings were sorted by ISI and by the perceptual outcome of the preceding trial (t–1 integration vs. t–1 segregation), and compared using permutation-based paired t-tests (10,000 permutations; FDR-corrected, q = 0.05). Confidence ratings were significantly higher following t–1 integration trials at short ISIs (21–63 ms; q < 0.05; Fig. 1D), whereas confidence was higher following t–1 segregation trials at long ISIs (104–125 ms; q < 0.05; Fig. 1D). No significant effects of prior perception emerged at the extremes of the temporal range (0 ms and 146 ms; q > 0.05; Fig. 1D) or near the point of subjective equality (83 ms; q > 0.05).

Next, we tested whether confidence scaled with perceptual-evidence strength, increasing as current evidence became stronger relative to each participant’s own temporal threshold (i.e., PSE). To this end, we modelled trial-by-trial subjective confidence reports using a hierarchical Bayesian ordered-logistic framework (see Methods). Confidence increased markedly with absolute perceptual evidence (β = 1.683, 95% HDIs [1.537, 1.826], Pr(β > 0) > .999; 95% HDIs [4.65, 6.21]; Fig. 1E). Here, absolute perceptual evidence indexed the magnitude of current stimulus evidence relative to each participant’s own PSE, irrespective of perceptual direction. The underlying signed evidence was increasingly negative as integration-favouring evidence became stronger within the participant-specific integration window, and increasingly positive as segregation-favouring evidence became stronger within the participant-specific segregation window. Thus, moving farther from the individual perceptual boundary (PSE) in either direction was associated with progressively higher confidence. Because perceptual-evidence strength was centred within participant (i.e., individual PSE), this effect indicates that confidence systematically increased with trial-to-trial variations in perceptual evidence within individuals, independently of stable between-participant differences in confidence-scale use. The strength of this confidence– evidence coupling also varied across participants (evidence slope SD = 0.622, 95% HDIs [0.512, 0.741]). In parallel, participants showed substantial individual variation in their general confidence tendency (intercept SD = 1.245, 95% HDIs [1.028, 1.457]). Thus, confidence systematically tracked individualized perceptual-evidence strength, while the hierarchical model separately captured individual differences in both the strength of this coupling and general confidence tendency. Consistent with the preceding model-free analysis, confidence was additionally modulated by the congruence between current perceptual evidence and recent perceptual experience. The perceptual history-congruence effect was positive (β = 0.243, 95% HDIs [0.186, 0.305], Pr(β > 0) > .999), indicating greater confidence when current evidence favoured the same perceptual interpretation reported on the preceding trial, whether integration or segregation. In contrast, there was no consistent difference in confidence between preceding integration and segregation at the perceptual boundary (β = 0.011, 95% HDIs [−0.060, 0.078]), indicating that perceptual history influenced confidence primarily as a function of its congruence with current perceptual evidence.

Together, these findings indicate that confidence systematically tracked the strength of individualized perceptual evidence and was further enhanced when current temporal evidence supported the same perceptual outcome as the previous trial, suggesting that recent perceptual states bias not only perceptual judgments but also their subjective certainty^61^.

### Resting-state oscillatory and aperiodic neural dynamics

Alongside the temporal integration task, we recorded resting-state EEG activity (eyes-closed, EC; eyes-open, EO) from 64 scalp electrodes. In order to examine how resting-state oscillatory and aperiodic neural dynamics jointly predict visual temporal processing, we applied the FOOOF algorithm^32^ to decompose each participant’s power spectrum into oscillatory and aperiodic components (Fig. 2A). Spectral parameterization provided excellent fits across participants and electrodes, with median R^2^ values of 0.9936 in EC and 0.9933 in EO (fitting error = 0.0327 and 0.0297, respectively). Following spectral parameterization, we quantified the prevalence of identified oscillatory peaks across participants and electrodes. Alpha peaks (8–13 Hz) were particularly robust, being identified in 99.4% and 98.1% of all participant–electrode spectra, estimated separately for each participant and scalp electrode, in EC and EO, respectively. As expected, peak identification outside the alpha range was more heterogeneous (see Methods).

**Figure 2.**
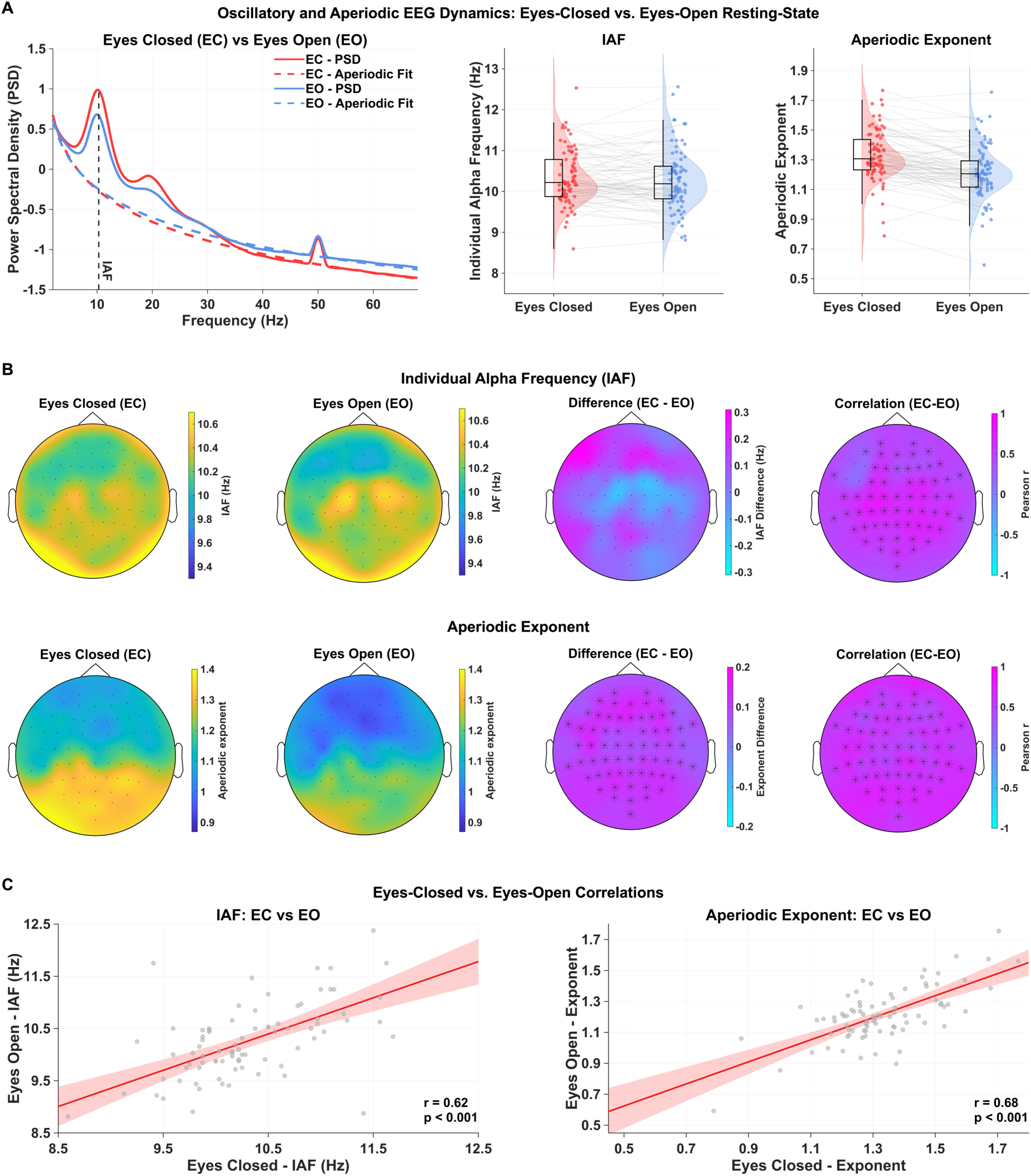
Resting-state oscillatory and aperiodic neural dynamics across eyes-closed and eyes-open conditions. (A) Group-averaged resting-state power spectra for eyes-closed (EC; red) and eyes-open (EO; blue) conditions. Solid lines show power spectral density (PSD), wereas dashed lines show the fitted aperiodic component. The vertical dashed line indicates the group-level individual alpha frequency (IAF). Adjacent violin plots show participant-level aperiodic-corrected IAF and aperiodic exponent for EC and EO resting-state conditions (averaged across posterior electrodes), with individual participants connected across conditions. (B) Scalp distributions of IAF (top row) and aperiodic exponent (bottom row). From left to right, topographical maps show EC, EO, the EC–EO difference, and the across-participant EC–EO cross-state correlation at each electrode. Difference maps represent the mean within-participant EC–EO difference, whereas correlation maps represent the channel-wise Pearson correlation between EC and EO values across participants. Asterisks indicate electrodes belonging to clusters identified by the corresponding cluster-permutation analysis (10,000 permutations; cluster-corrected p < 0.05). No significant EC–EO difference cluster was identified for IAF, whereas the aperiodic exponent was higher during EC than EO across a widespread scalp cluster. Both IAF and aperiodic exponent showed widespread positive cross-state correlations. (C) Participant-level EC–EO relationships for posterior-ROI IAF (left) and aperiodic exponent (right). Points represent individual participants; solid lines indicate the linear regression fit and shaded regions the 95% confidence interval. EC and EO values were positively correlated for both IAF (r = 0.62) and aperiodic exponent (r = 0.68). Statistical inference was based on cluster-based permutation testing at the scalp level.

We then examined the relationship between EC and EO resting-state activity across our two primary neural measures of interest: individual alpha frequency (IAF) and the aperiodic exponent. IAF was quantified from the aperiodic-corrected spectrum as the frequency exhibiting maximum power within the alpha range (8–13 Hz). To validate this measure against an independently derived spectral parameter, we compared aperiodic-corrected IAF estimates with with the centre frequency peaks independently identified by FOOOF. The two estimates showed very high agreement across participants in both EC (*r* = 0.975, p < 0.001) and EO (*r* = 0.976, p < 0.001) resting conditions. IAF (aperiodic-corrected) did not differ between EC and EO at the scalp level (Fig. 2B), with no clusters identified in the paired cluster-permutation analysis (10,000 permutations, p-values > 0.05; Fig. 2B). Participants’ IAF estimates were strongly related across resting states: EC and EO IAFs showed a positive cross-state association across a widespread cluster of electrodes (*p* < 0.001; Fig. 2B/C). In contrast, while showing strong positive cross-state associations across the scalp (*p* < 0.001; Fig. 2B/C), the aperiodic exponent was significantly higher across widespread scalp regions in the EC condition compared to EO (*p* < 0.001; Fig. 2B).

Together, these results indicate a substantial intra-individual correlation between resting states for both IAF and aperiodic activity. Nonetheless, the aperiodic spectrum was significantly steeper in the EC state, whereas IAF exhibited no significant modulations, consistently with the view that aperiodic broadband neural activity reflects state-dependence mechanisms^27,29,32^, while alpha speed represents a relatively stable neural trait^51,52,56,57^.

### Alpha frequency and aperiodic neural dynamics predict temporal integration

Next, we directly tested the oscillatory sampling hypothesis, correlating resting-state IAF (aperiodic-corrected) with behavioral measures of temporal integration (PSE and psychometric slope) using Pearson correlations with cluster-based correction (10,000 permutations). All correlation analyses were performed separately for eyes-closed (EC) and eyes-open (EO) resting-state measures. Faster IAFs, particularly across posterior–central scalp regions, were associated with lower temporal thresholds (PSE) in both EC (p = 0.0017) and EO (p = 0.0001) conditions (Fig. 3A), supporting the notion that faster resting-state alpha rhythms predict narrower integration windows. By contrast, IAF was not significantly correlated with psychometric slope in either condition (p > 0.05; Fig. 3B). We then examined the contribution of aperiodic neural activity. Higher aperiodic exponents were significantly associated with steeper psychometric slopes in both EC (p = 0.0014) and EO (p = 0.0001) conditions (Fig. 3D), indicating that a steeper (less flat) spectrum was linked to greater precision in temporal binding judgments. By contrast, aperiodic exponent was not significantly associated with temporal thresholds (PSE) in either EC or EO conditions (p > 0.05; Fig. 3C). Importantly, neither resting-state IAF nor aperiodic exponent was associated with the estimated rate of stimulus-independent responses (lapse rate), with no significant scalp-level associations observed in either EC or EO recordings (cluster-corrected p-values > .05). Thus, the observed relationships between resting-state spectral measures and psychometric parameters were unlikely to be driven by individual differences in stimulus-independent responding.

**Figure 3.**
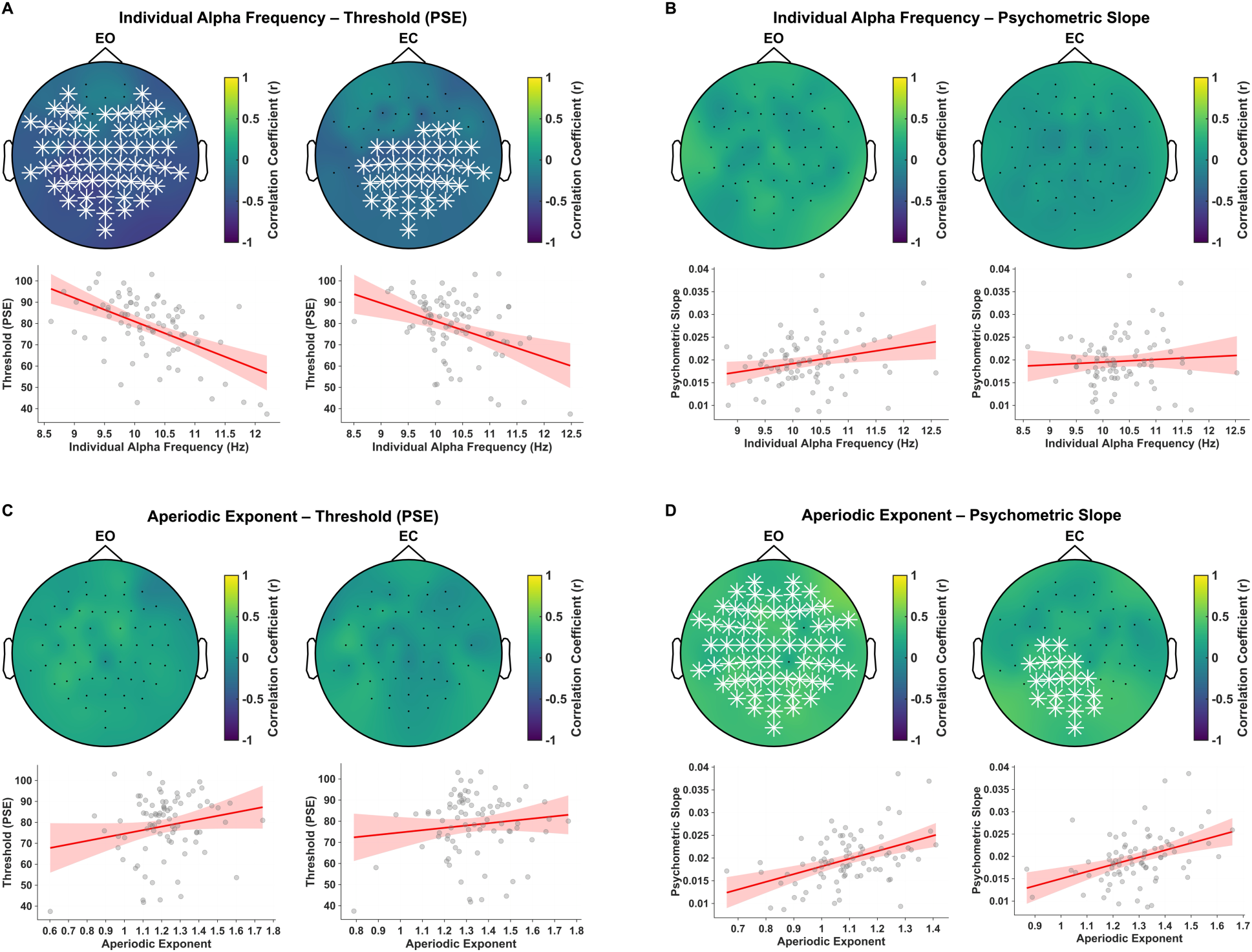
Resting-state oscillatory and aperiodic neural dynamics predict visual temporal integration performance. (A) Topographical maps show the scalp distribution of correlations between individual alpha frequency (IAF) and temporal threshold of the psychometric function (PSE) during eyes-open (EO) and eyes-closed (EC) resting-state conditions. Faster IAFs sites were significantly associated with lower temporal thresholds in both EO (p = 0.0001) and EC (p = 0.0017) conditions (cluster-corrected), indicating finer temporal resolution and narrower integration windows. (B) Correlations between IAF and psychometric slope revealed no significant effects in either resting-state condition (p > 0.05), suggesting that alpha frequency primarily relates to temporal binding boundaries rather than perceptual precision. (C) Topographies and scatterplots revealed no significant associations between aperiodic exponent and temporal threshold in either EO or EC conditions (p > 0.05). (D) Higher aperiodic exponents were positively correlated with steeper psychometric slopes in both EO (p = 0.0001) and EC (p = 0.0014) conditions (cluster-corrected), indicating that steeper aperiodic spectra were associated with greater precision in temporal binding judgments. Together, these results reveal complementary contributions of oscillatory and aperiodic neural dynamics to visual temporal processing, with faster alpha rhythms associated with narrower temporal integration windows and steeper aperiodic spectra associated with steeper psychometric functions. Statistical inference was based on cluster-based permutation testing at the scalp level. Scatterplots are shown for visualization, and neural measures were averaged across electrodes belonging to the significant cluster when present, and across a predefined posterior-central electrode set for analyses without a significant cluster.

To assess the spectral specificity of the associations of resting-state IAF and aperiodic exponent with temporal perception, we performed analogous control analyses using aperiodic-corrected peak-frequency estimates outside the alpha range. As expected, while alpha peaks showed high identifiability across participants and electrodes, the prevalence of peaks identified during spectral parametrization was more heterogeneous and less reliable across the other frequency bands (see Methods). Despite this variability, analogous aperiodic-corrected peak-frequency estimates were derived for the delta (1–4 Hz), theta (4–7 Hz), low-beta (15–30 Hz), high-beta (30–45 Hz), and gamma (45–70 Hz) ranges. None of these non-alpha frequency measures showed a significant association with either PSE or psychometric slope in EC or EO recordings (p-values > 0.05; see Methods), supporting the frequency specificity of the relationship between IAF and temporal integration. Similarly, given previous evidence linking resting-state oscillatory power to temporal integration across multiple timescales^62–64^, we additionally examined aperiodic-corrected oscillatory power in the delta (1–4 Hz), theta (4–7 Hz), alpha (8–13 Hz), low-beta (15–30 Hz), high-beta (30–45 Hz), and gamma (45–70 Hz) ranges. None of these power measures showed a significant association with either PSE or psychometric slope in EC or EO recordings (p-values > 0.05).

Together, these findings corroborate previous evidence that visual temporal acuity emerges from the joint influence of oscillatory alpha dynamics and aperiodic neural mechanisms^27,29,33,34^.

### IAF predicts temporal integration beyond stimulus – IAF frequency matching

Because the sustained stimulus sequence alternated continuously between the two Gabor orientations, some stimulus durations produced temporal structure within the alpha-frequency range. This is potentially relevant because rhythmic visual stimulation can elicit strong stimulus-locked responses in the alpha band, and its behavioural effects may depend on the correspondence between the stimulation frequency and an individual’s intrinsic alpha frequency^56,58,65–68^. We therefore tested whether the observed association between resting IAF and temporal-integration PSE could be accounted for by the proximity between IAF and the alpha-range frequencies present in the stimulus sequence. Specifically, using hierarchical Bayesian psychometric modeling (see Methods), we compared a model in which resting IAF (averaged across posterior electrodes) predicted participant-level PSE (IAF-only model) with an extended model that additionally incorporated stimulus frequency–IAF distance as a predictor of the estimated PSE. This distance term was defined for the three stimulus-duration conditions whose corresponding alternation rates fell within the alpha range (8–13 Hz): 12 frames (83.33 ms; 12 Hz), 15 frames (104.17 ms; 9.6 Hz), and 18 frames (125 ms; 8 Hz).

In the IAF-only model, higher resting IAF was associated with lower PSEs (i.e., narrower temporal integration windows) in both resting-state conditions. For EC, a 1-SD increase in IAF was associated with a 4.36-ms decrease in PSE (β_IAF_ = −4.36 ms/SD, 95% HDIs [−7.22, −1.51], Pr[β_IAF_ < 0] = .998). Similarly, the corresponding association was larger for EO (β_IAF_ = −7.34 ms/SD, 95% HDIs [−9.73, −4.63], Pr[β_IAF_ < 0] > .999). Critically, these associations were essentially unchanged when stimulus frequency–IAF distance was included in the model (Fig. S1). For EC, β_IAF_ was −4.30 ms/SD (95% HDIs [−7.22, −1.48], Pr[β_IAF_ < 0] = .998), corresponding to only a 1.43% reduction in the absolute magnitude of the IAF effect relative to the IAF-only model. For EO, β_IAF_ was −7.40 ms/SD (95% HDIs [−10.01, −4.85], Pr[β_IAF_ < 0] > .999), with the absolute magnitude of the association instead increasing slightly by 0.76%. By contrast, there was no clear evidence that relative stimulus frequency–IAF distance independently shifted the temporal-integration threshold. The posterior distribution of β_distance_ was centered close to zero for both EC (β_distance_ = 0.28 ms/Hz, 95% HDIs [−1.26, 1.74], Pr[βdistance > 0] = .642) and EO (β_distance_ = −0.16 ms/Hz, 95% HDIs [−1.59, 1.35], Pr[β_distance_ > 0] = .416; Fig. S1). Importantly, including stimulus frequency–IAF distance did not improve out-of-sample predictive accuracy. Model comparisons indicated that the two models were predictively indistinguishable for EC (ΔELPD = 0.45, SE = 1.06), whereas for EO the IAF plus distance model showed lower predictive accuracy than the IAF-only model (ΔELPD = 2.86, SE = 0.79).

Together, these analyses indicate that the relationship between faster resting IAF and lower temporal integration thresholds was robust to individual differences in the correspondence between endogenous alpha frequency and the alpha-range temporal structure of the stimulus. Although the present data cannot directly determine the neural response elicited by the stimulus sequence in the absence of task-related EEG activity, the resting-state IAF–PSE association was not accounted for by stimulus frequency–IAF distance.

### Alpha rhythm and aperiodic exponent predict the weighting of prior experience

We next examined whether resting-state EEG activity predicts the strength of serial dependence during temporal binding of visual information. For each participant, we quantified serial dependence from the posterior participant-specific PSEs for trials following t–1 integration and t– 1 segregation, computing their difference as ΔPSE = PSE t–1 integration – PSE t–1 segregation. This index captures how strongly the percept from the previous trial biases current temporal judgments. Correlating this behavioral index with resting-state EEG measures (IAF and aperiodic exponent) revealed distinct neural signatures of serial dependence. Faster IAFs, particularly over posterior electrodes, were associated with lower ΔPSE values in the EO condition (p = 0.0430; Fig. 4A), indicating that individuals with faster resting-state alpha rhythms (i.e., finer temporal resolution) were less influenced by prior experience, relying more on current sensory evidence. This relationship was absent in the EC condition (p > 0.05). Conversely, lower aperiodic exponents were linked to higher ΔPSE values in both EO (p = 0.0006) and EC (p = 0.0045) conditions (Fig. 4B), suggesting that individuals with flatter resting-state aperiodic spectra may rely more heavily on prior perceptual outcomes to stabilize perceptual decisions under uncertainty.

**Figure 4.**
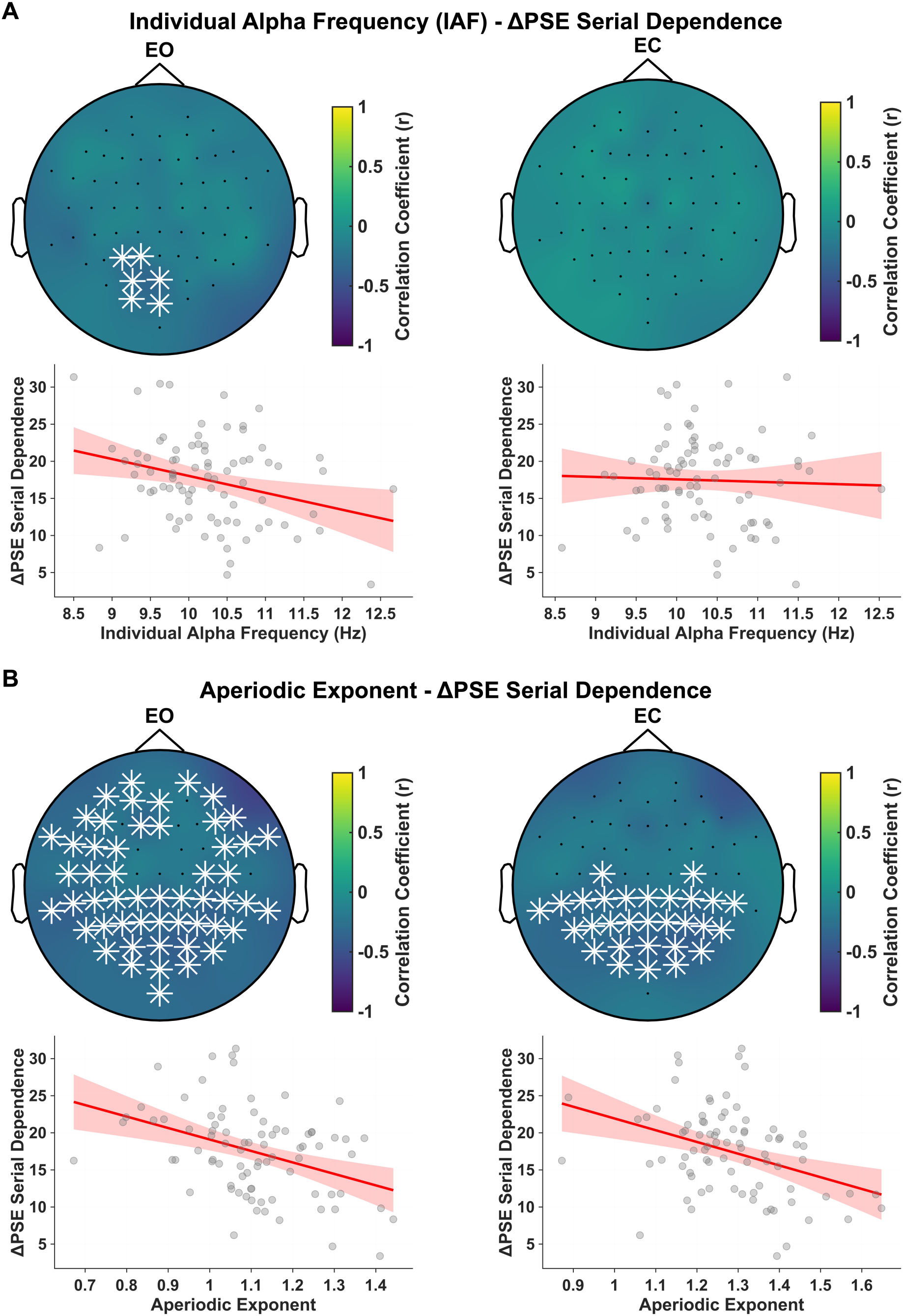
Resting-state neural dynamics predict individual differences in serial dependence. (A) Topographies show correlations between individual alpha frequency (IAF) and serial dependence strength (ΔPSE = PSE t–1 integration – PSE t–1 segregation) for eyes-open (EO) and eyes-closed (EC) conditions. Faster IAFs over posterior electrodes were associated with lower ΔPSE in the EO condition (p = 0.0430), indicating that individuals with faster alpha rhythms (i.e., higher temporal acuity) relied less on prior perceptual outcomes. No significant effects were observed in the EC condition (p > 0.05). (B) Correlations between the aperiodic exponent and ΔPSE revealed that flatter spectra (lower exponents) were linked to higher ΔPSE values in both EO (p = 0.0006) and EC (p = 0.0045) conditions, suggesting greater dependence on perceptual history to stabilize perceptual decisions under uncertainty. Together, these findings indicate that resting-state oscillatory and aperiodic features both predict the magnitude of reliance on prior experience, with faster alpha oscillations and steeper neural spectra associated with reduced serial dependence. Scatterplots are shown for visualization, and neural measures were averaged across electrodes belonging to the significant cluster when present, and across a predefined posterior-central electrode set for analyses without a significant cluster.

Finally, given that both IAF and aperiodic exponent predicted serial dependence strength (ΔPSE) in the EO resting-state condition, we performed reciprocal mediation analyses (see Methods). IAF and aperiodic exponent were not significantly associated with one another (β = −0.041, p = .794). Accordingly, there was no evidence that the relationship between IAF and ΔPSE was mediated by aperiodic exponent (indirect effect: β = 0.016, 95% bootstrap CI [−0.099, 0.156], p = .797), nor that the relationship between aperiodic exponent and ΔPSE was mediated by IAF (β = 0.012, 95% bootstrap CI [−0.087, 0.126], p = .812). Instead, both neural measures remained independently associated with serial dependence when entered simultaneously: faster IAF predicted lower ΔPSE (β = −0.308, 95% CI [−0.490, −0.109], p = .002), as did higher aperiodic exponent (β = −0.402, 95% CI [−0.594, −0.211], p < .001).

Together, these findings indicate that resting-state oscillatory and aperiodic features make separable contributions to individual differences in reliance on prior experience, with faster alpha oscillations and steeper aperiodic spectra each associated with reduced dependence on the immediate perceptual past.

### Neural predictors of subjective confidence in temporal binding

Finally, we tested whether resting-state EEG measures predicted subjective confidence in temporal binding judgments. We first examined participants’ mean confidence ratings. Faster IAFs were associated with higher mean confidence across widespread scalp regions in both the EO (p = 0.0001) and EC (p = 0.0011) conditions (Fig. 5A), indicating that individuals with faster resting-state alpha rhythms tended to report greater subjective confidence in their perceptual judgments. On the other hand, no significant relationships were observed between the aperiodic exponent and mean confidence (p > 0.05). To assess the spectral specificity of the relationship between IAF and confidence, we performed analogous control analyses using aperiodic-corrected peak-frequency estimates in the delta (1–4 Hz), theta (4–7 Hz), low-beta (15–30 Hz), high-beta (30–45 Hz), and gamma (45–70 Hz) ranges. None of these non-alpha frequency measures showed a cluster-based association with mean confidence in either EC or EO recordings (p-values > 0.05). We additionally examined aperiodic-corrected oscillatory power across the delta, theta, alpha, low-beta, high-beta, and gamma ranges. None of these power measures was associated with mean confidence in either resting-state condition (p-values > 0.05), indicating that the relationship with confidence was specific to IAF rather than oscillatory power.

**Figure 5.**
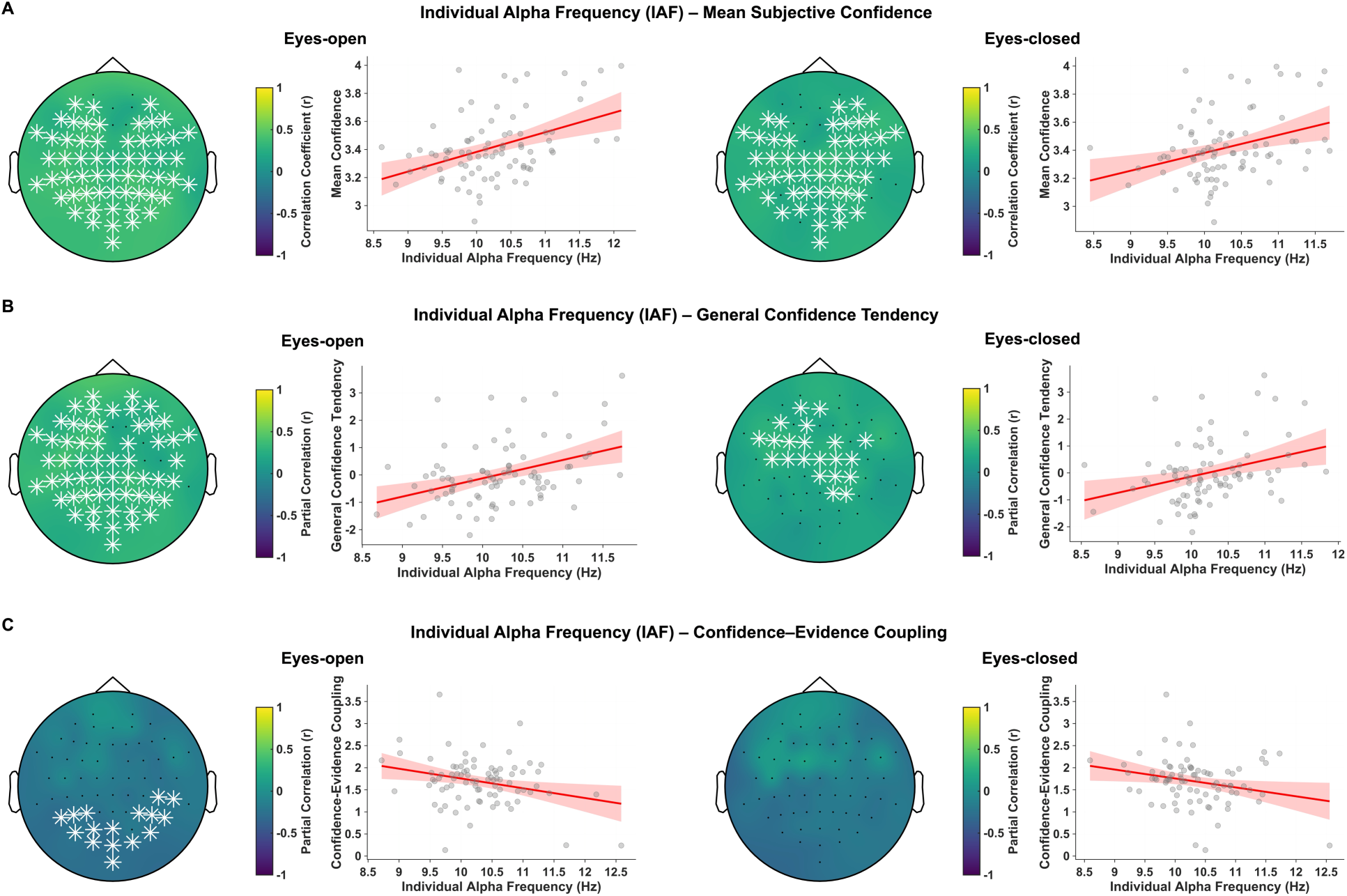
Resting-state individual alpha frequency predicts distinct components of subjective confidence. (A) Topographies show scalp-level correlations between individual alpha frequency (IAF) and mean subjective confidence for eyes-open (EO) and eyes-closed (EC) resting-state conditions. Faster IAFs were associated with higher mean confidence across widespread scalp regions in both EO (p = 0.0001) and EC (p = 0.0011) conditions, indicating that individuals with faster alpha rhythms tended to report greater subjective confidence in their temporal binding judgments. (B) To decompose this association, we examined the participant-specific confidence parameters estimated by the hierarchical ordered-logistic model. General confidence tendency corresponded to the participant-specific intercept and indexed the general tendency to report higher versus lower confidence, conditional on the remaining model predictors. Partial-correlation analyses controlling for confidence–evidence coupling revealed that faster IAFs were associated with higher general confidence tendency in both EO (p = 0.0004) and EC (p = 0.0194) conditions. (C) Confidence–evidence coupling corresponded to the participant-specific absolute-evidence slope and indexed how strongly confidence tracked trial-to-trial variations in individualized perceptual-evidence strength, defined relative to each participant’s perceptual boundary (PSE). Partial correlations controlling for general confidence tendency showed that faster IAFs were associated with weaker confidence–evidence coupling over posterior electrodes in EO (p = 0.0258), whereas this relationship did not reach the corrected criterion in EC (p = 0.0627). No relationships were observed between the aperiodic exponent and mean confidence, general confidence tendency, or confidence–evidence coupling in either resting-state condition (p-values > 0.05). Together, these findings indicate that IAF predicts dissociable components of subjective confidence: faster alpha rhythms are associated with a greater general tendency toward higher confidence, while also showing a weaker trial-by-trial coupling between confidence and perceptual-evidence strength. White stars indicate electrodes belonging to clusters surviving cluster-based correction. Scatterplots are shown for visualization, and neural measures were averaged across electrodes belonging to the significant cluster when present, and across a predefined posterior-central electrode set for analyses without a significant cluster. For partial-correlation scatterplots, covariate-adjusted values were shifted back to the original parameter mean for visualization, without altering the underlying partial correlation.

We next asked which component of confidence accounted for the association with IAF by leveraging the participant-specific parameters estimated by the hierarchical ordered-logistic confidence model (see above and Methods). Specifically, we distinguished general confidence tendency, corresponding to the participant-specific intercept and reflecting the general tendency to report higher versus lower confidence conditional on the remaining model predictors, from confidence–evidence coupling, corresponding to the participant-specific absolute-evidence slope and reflecting how strongly confidence tracked trial-to-trial variations in individualized perceptual-evidence strength (i.e., distance from PSE). To isolate the unique neural association of each confidence component, we performed scalp-level partial-correlation analyses in which the relationship between IAF and one confidence parameter was estimated while controlling for individual differences in the other parameter. Thus, the analysis of general confidence tendency tested its association with IAF after removing variance shared with confidence–evidence coupling, whereas the complementary analysis of confidence–evidence coupling controlled for general confidence tendency. Scalp-level partial-correlation analyses revealed that faster IAFs were associated with higher general confidence tendency in both EO (p = 0.0004) and EC (p = 0.0194) conditions (Fig. 5B), independently of individual differences in confidence–evidence coupling. By contrast, faster IAFs were associated with weaker confidence–evidence coupling over posterior electrodes in the EO condition (p = 0.0258; Fig. 5C), independently of general confidence tendency, whereas this relationship did not reach the corrected significance criterion in EC (p = 0.0627). No relationships were observed between the aperiodic exponent and general confidence tendency, or confidence–evidence coupling in either resting-state condition (all p-values > 0.05). Together, these findings indicate that resting-state IAF predicts distinct components of subjective confidence: faster alpha rhythms are associated with a greater general tendency toward higher confidence, while they are also associated with a weaker trial-by-trial coupling between confidence and perceptual-evidence strength, reflecting a higher general level of subjective certainty while reducing the relative contribution of moment-to-moment evidence fluctuations to confidence formation.

## Discussion

The perceptual system segments the continuous flow of sensory inputs into discrete events to construct a coherent experience of an otherwise chaotic world. This temporal parsing has long been linked to the intrinsic speed of alpha-band oscillations^2^ and more recently to the scale-free, aperiodic organization of neural activity^27,29,34^. However, recent studies have reported contradictory findings that challenge a direct link between alpha oscillations and individual differences in perceptual resolution^6^. Here, by combining a novel continuous temporal integration paradigm with resting-state EEG, we replicate the classical two-flash findings while overcoming their key limitations that can confound genuine perceptual binding from failure to fully process both stimuli. We replicated the finding that resting-state oscillatory and aperiodic activity jointly predict the threshold and slope of the psychophysical curve^5,27^, now using a sustained visual stream, as well as linking it for the first time to previous perception effects and confidence measures. Our results reveal that visual temporal integration may arise from a multilevel process governed by the interplay of rhythmic and aperiodic neural activity, perceptual history, and subjective confidence, moving beyond a simple account of the role of alpha oscillations in temporal integration.

### Alpha rhythm and aperiodic exponent jointly predict visual temporal performance

First, our findings provide support for, but also a more complex understanding of the link between brain rhythms and visual perception. Individuals with faster resting-state alpha rhythms did indeed show faster temporal integration thresholds, in line with previous findings using brief flashes^2–4,9,10,27,29,69^. However, the speed of alpha frequency did not predict the steepness of the psychometric function (slope), suggesting that resting-state oscillatory activity alone does not account for individual differences in visual temporal perception^27,29,34,70,71^.

Our findings revealed that this aspect of temporal processing was captured by the aperiodic component of resting-state neural activity^27,29,34^. Higher aperiodic exponents were associated with steeper psychometric slopes, indicating that a steeper neural spectrum was linked to greater sensitivity to temporal evidence. Importantly, because both neural measures were acquired at rest, these relationships should be interpreted as associations between relatively stable individual characteristics and behavioral performance. Within this framework, our findings are consistent with the view that the broadband cortical state in which endogenous neural activity unfolds may predict stable individual differences in temporal binding performance and contribute to how sensory information is bound over time. From this perspective, flatter neural spectra have been proposed to characterize neural states with reduced temporal coordination, whereas steeper spectra may reflect stronger inhibitory constraints and more temporally organized neural activity^29,31,46–48^. In this regard, both steeper resting-state and task-related neural spectra have been associated with reduced neural variability and increased signal-to-noise ratio, potentially supporting more reliable sensory processing^27,29,72^. Thus, the association observed here between steeper resting-state spectra and steeper psychometric slopes is consistent with the possibility that cortical states characterized by lower neural variability at rest may predict more reliable perceptual representations and higher sensitivity to temporal evidence. These findings further emphasize and extend the potential relevance of the aperiodic structure of electrophysiological activity for cognition and perception, which has been associated with multiple domains including processing speed^39,54^, working memory^73^, emotional processing and arousal^38,74^, as well as aging^29,36,46,72^, and a broad range of psychopathological conditions^29,36^.

Importantly, these findings support complementary roles for resting-state oscillatory and aperiodic neural dynamics in visual temporal perception. Alpha frequency may constrain the temporal scale at which sensory information is sampled and integrated, whereas the broadband aperiodic cortical state may influence how reliably and precisely information is processed within these oscillatory sampling windows through influences on neuronal gain and processing variability^2,27,29,34^. At a mechanistic level, it has been proposed that lower cortical variability allows rhythmic fluctuations to exert a stronger influence on sensory processing^27,50^. Accordingly, our findings suggest that individual differences in perceptual experience may depend jointly on the temporal organization indexed by resting-state oscillatory activity and on the broadband cortical state in which those oscillatory processes unfold.

### Alpha rhythm and aperiodic exponent predict the weighting of prior experience

The current findings explain in more detail how resting state EEG can predict individual performance in temporal perception tasks. Our findings reveal that visual temporal integration judgments are strongly perceptual history-dependent and shaped by both resting-state alpha rhythmicity and aperiodic neural activity. When we separately examined temporal integration thresholds for trials preceded by an integration response and those preceded by a segregation response, we found that temporal binding judgments were biased toward the previous perceptual outcome. Specifically, a previous integration response increased the likelihood of perceiving a subsequent stimulus as a single unified event, and vice-versa for segregation responses. These results demonstrate that recent perceptual experience flexibly biases the temporal binding window, consistent with recent evidence showing that temporal perception reflects a dynamic interplay between the temporal structure of sensory evidence and perceptual history^15–17^.

Recent work has suggested that alpha oscillations may also support serial dependence processes^14,75^. In this study, faster resting-state alpha rhythms and higher aperiodic exponents (i.e., steeper neural spectra) were associated with a lower magnitude of serial dependence effect, indicating reduced influence of prior perceptual outcomes and a greater reliance on current sensory evidence, potentially due to increased perceptual precision^18,19^. Conversely, individuals characterized by slower alpha rhythms and lower aperiodic exponents showed stronger serial dependence. This pattern of results may suggest that resting-state oscillatory and aperiodic neural dynamics predict individual differences in susceptibility to prior perceptual history, whereby slower oscillatory sampling and greater neural variability may index a greater tendency to weight prior perceptual information when current sensory evidence is more uncertain^14^. Importantly, we performed mediation analyses to examine whether these associations reflected interdependent relationships or, instead, separable contributions to individual differences in reliance on prior experience. Both resting-state alpha frequency and aperiodic slope remained independently associated with the magnitude of the serial dependence effect, suggesting that oscillatory and aperiodic features may make separable contributions. However, given the resting-state nature of the current electrophysiological data, whether oscillatory and aperiodic neural activity dynamically regulate the weighting of prior experience relative to current sensory information, in a manner resembling the associations observed here with resting-state measures, remains to be established.

### Resting-state alpha frequency as a neural predictor of subjective confidence

After each temporal binding judgment, participants rated their confidence, which mirrored the clarity of temporal evidence: confidence was lowest near the temporal integration threshold and highest at short and long inter-stimulus intervals, where evidence more clearly supported either integration or segregation. Additionally, subjective confidence was inversely related to individual temporal thresholds, indicating that observers with narrower integration windows reported higher overall confidence in their perceptual judgments. Trial-level hierarchical modelling showed that confidence increased with the absolute strength of individualized perceptual evidence (i.e., distance from PSE). This pattern is consistent with evidence-strength accounts of perceptual confidence, in which subjective certainty increases as internal evidence moves farther from the decision boundary^24–26^. Crucially, confidence was also modulated by the congruence between prior perceptual history and current temporal evidence. Confidence ratings were significantly higher following integration judgments at short inter-stimulus intervals and following segregation judgments at long intervals. Thus, confidence was enhanced when current sensory evidence favoured the same perceptual outcome as the preceding trial, indicating that recent perceptual states influence not only perceptual judgments but also the subjective certainty assigned to them^61^. This result converges with evidence that prior information and sequential dependencies can systematically shape confidence judgments, particularly when prior and current information are congruent^76,77^.

Finally, we tested whether resting-state neural dynamics predicted subjective confidence in temporal binding judgments. Trial-level modelling of confidence revealed that faster IAF was associated with a greater general confidence tendency, suggesting that a relatively stable feature of intrinsic neural activity such as resting-state alpha frequency^51,52,56^ covaries with an individual’s general propensity to experience higher subjective confidence. This association is consistent with a trait-level interpretation, as both resting-state IAF and the general confidence parameter capture stable between-subject differences. At the same time, faster IAF was independently associated with weaker confidence–evidence coupling, indicating that confidence in individuals with faster resting-state alpha rhythms was less strongly modulated by trial-to-trial variations in temporal-evidence strength. Thus, faster alpha rhythms predicted separable components of subjective confidence: a higher general level of subjective certainty and a reduced dependence of confidence on momentary fluctuations in perceptual evidence. One possible interpretation is that individuals characterized by faster intrinsic alpha rhythms maintain a higher and more stable general confidence level, such that confidence varies less strongly with fluctuations in the strength of current sensory evidence. However, because IAF was measured at rest, this interpretation should be regarded as a trait-level association rather than evidence that alpha frequency dynamically regulates confidence during individual trials.

### Oscillatory and aperiodic dynamics shape visual performance beyond temporal integration

Overall, the alpha rhythm was distinctly associated with multiple facets of visual temporal processing, from temporal thresholds and serial dependence to subjective confidence. Beyond the resting-state associations observed here, one potential explanation for this multilevel influence of alpha oscillatory speed on temporal binding is that faster alpha rhythms operating within a steeper aperiodic neural background characterized by lower neural variability may enhance temporal resolution through more efficient sensory evidence accumulation^2,5,18,19,27,29,33,34,51,58,69,78^. Recent evidence suggests that faster alpha rhythms, measured both at rest and during task performance, predict greater perceptual resolution and may promote it by generating more sampling frames per time unit, thereby increasing the amount of accumulated sensory evidence^18,19,34^. Consequently, more accurate and higher-rate sampling of visual input may yield a more faithful representation of external events, resulting in higher temporal precision, reduced reliance on prior perceptual experience, and greater subjective confidence in perceptual decisions, as observed in the current study. Together, these results support and expand on our recently proposed framework linking alpha oscillatory speed and aperiodic neural dynamics in perceptual integration^5,27,33,34^, showing also a link to decisional and metacognitive facets. By linking rhythmic sampling with the stability of the underlying neural background, this framework highlights that temporal perception emerges from the coordinated interplay between oscillatory precision, the steepness of the neural spectrum, and the brain’s predictive use of recent perceptual history. This account provides a broader model for how rhythmic and aperiodic neural dynamics jointly sustain the continuity of conscious visual experience across time.

### Limitations and future directions

While our findings reveal robust associations between resting-state dynamics and visual temporal performance, they should be interpreted as relationships between stable neural features and behavior, and cannot be directly extended to neural dynamics occurring during task performance. Nevertheless, the finding that resting-state alpha frequency and the aperiodic exponent showed robust associations with subsequent perceptual performance across both resting-state conditions is consistent with prior evidence that resting-state neural indices are relatively stable within individuals^41,51–54,56,57^ and predict interindividual differences in cognition and perception^27,41,51^. However, whether oscillatory and aperiodic neural activity dynamically regulate the different aspects of visual temporal performance identified here, in a manner resembling their resting-state associations, remains to be established. Future studies employing task-related M/EEG designs would provide important complementary information by allowing a direct characterization of how pre-stimulus and ongoing neural dynamics relate to visual temporal perception. Indeed, moment-to-moment perceptual sensitivity can vary with oscillatory phase and power, consistent with alternating periods of relatively greater and lower excitability across the alpha cycle^2,3,79^, and these oscillatory effects may depend on the ongoing aperiodic cortical state^27,37,47,50^, such that phase-dependent fluctuations and broadband cortical state may jointly shape perception.

Relatedly, the present design does not allow us to determine whether these psychometric parameters primarily reflect sensory or decisional processes. Indeed, temporal integration thresholds may reflect differences in temporal sensitivity or biases toward temporal integration versus segregation, whereas the variability in the steepness of the perceptual transition may arise from both sensory encoding and decisional processes^59,60^. Future implementations of the present paradigm could be adapted to disentangle these contributions. Signal-detection and evidence-accumulation approaches could help dissociate sensory and decisional contributions to perceptual performance and their associations with neural measures, for example by incorporating signal-present and signal-absent reference conditions and allowing responses during stimulus presentation so that reaction times meaningfully reflect the decision process.

Additionally, our findings also revealed how eyes-closed (EC) and eyes-open (EO) resting-state activity are related and may differentially predict visual temporal performance. EC and EO are considered related resting conditions in which overlapping neural features are expressed under different state constraints^38,80^. Our findings suggest that IAF is substantially preserved across EC and EO, consistent with previous work characterizing alpha frequency as a relatively stable neurophysiological trait^52,56,57^. At the same time, the aperiodic exponent showed cross-state correspondence within individuals together with systematic state dependence at the group level, consistent with evidence that aperiodic neural activity captures stable interindividual variability while remaining sensitive to changes in cortical state^27,29,32,41^. Relatedly, some brain–behavior associations were more evident in EO than EC. In line with previous findings^27^, one possible explanation is that EO provides a resting-state configuration more proximal to the visually engaged conditions under which the temporal task was performed, and this greater functional similarity may make EO measures particularly sensitive to individual differences relevant for visual processing. Future studies combining resting-state and task-related neural measures could help disentangle how stable neural traits and moment-to-moment fluctuations in neural dynamics relate to different stages and dimensions of visual temporal perception.

## Methods

### Participants

A total of 83 volunteers (46 females, mean age = 23.4 years, SD = 2.58) participated in the experiment and received compensation. Participants presented normal or corrected-to-normal vision. Exclusion criteria were self-reported neurological and attention disorders, epilepsy, and photosensitivity. The sample size was determined a priori based on a recent meta-analysis of 27 studies examining the relationship between individual alpha frequency and temporal resolution^2^. The meta-analysis reported a pooled population correlation of r = 0.394 (95% CI [0.332, 0.465]; SE = 0.034). Using this estimate as the expected effect size, an a priori power analysis (two-tailed, α = 0.05) indicated that a minimum of N = 49 participants would be required to achieve 80% power to detect r = 0.394, and N = 69 for the lower bound of the confidence interval (r = 0.332). To ensure sufficient power to detect effects within this plausible range, we recruited 83 participants, which provided approximately 96% power for r = 0.394. Power analyses were conducted using G*Power 3.1^81^. The experiment was conducted in accordance with the Declaration of Helsinki and approved by the New York University Abu Dhabi Human Research Protection Program Internal Review Board (IRB).

### Stimuli and Experimental Design

We designed a novel visual temporal integration task to measure individual integration performance while avoiding confounds associated with brief, discrete two-flash paradigms. Stimuli were generated and presented in MATLAB (MathWorks) using *Psychtoolbox* on a gamma-corrected monitor (refresh rate: 144 Hz). Observers viewed the display binocularly from a distance of 70 cm in a dimly lit room. On each trial, two Gabor patches were presented at the center of the screen, alternating over ∼1 s in the same spatial location. Each Gabor had a low spatial frequency (1.94 cycles/degree; λ = 0.52°) and a Gaussian envelope (σ = 0.62°), presented at full contrast, with orientations of +45° (right-tilted) and −45° (left-tilted). Each Gabor was rendered with a circular aperture and converted to a texture. These two orthogonal Gabors alternated rapidly to manipulate temporal perception. The key manipulation was the duration of each Gabor presentation (i.e., the inter-stimulus interval, ISI) rather than the interval between discrete events. The Gabors alternated in counterphase with an ISI ranging from 0 to 146 ms in 21-ms steps. Each ISI level was repeated 30 times, yielding 240 experimental trials presented in random order. For ISI = 0, a “fully fused” control was shown by presenting a precomputed fused texture with slight spatial jitter to prevent adaptation. The order of presentation (which orientation appeared first) was randomized across trials. This manipulation produced a continuous visual stream in which short presentation durations elicited the illusory percept of a single fused plaid (integration), whereas longer durations induced the perception of two alternating gratings (segregation). Because the stimuli had distinct orientations but occupied the same retinal location within an uninterrupted sequence, this paradigm dissociated genuine temporal binding from simple detection failures. At the beginning of the experimental session, resting-state EEG activity was recorded while participants relaxed for 3 minutes with eyes closed (EC condition) and 3 minutes with eyes open (EO condition). Before the task, participants received on-screen instructions and completed a short practice block to familiarize with the procedure. Each trial began with a central fixation cross (500 ms), followed by the ∼1 s alternation sequence. Immediately after stimulus offset, participants made a two-alternative forced choice indicating their percept: fused/single pattern (key F) or separate/alternating patterns (key S). They then rated their confidence on a four-point scale (1 = not confident at all, 2 = not very confident, 3 = quite confident, 4 = very confident). A short break was provided halfway through the session.

## Data Analysis

### Behavioral analyses

#### Visual temporal performance: Hierarchical Bayesian psychometric modelling

Temporal integration performance was analyzed using hierarchical Bayesian logistic psychometric modelling. The number of “separate” responses was modelled using a Binomial likelihood, with the probability of reporting the two stimuli as separate described as a logistic function of ISI. The psychometric function included three principal participant-level components: the Point of Subjective Equality (PSE), the psychometric slope, and the response asymptotes. The PSE corresponded to the ISI at which fused and separate percepts were equally likely. Lower PSE values indicated a greater tendency toward segregation and therefore a narrower temporal integration window, whereas higher PSE values indicated a greater tendency to integrate the two stimuli. The psychometric slope quantified how abruptly and consistently perceptual reports transitioned from integration to segregation across ISIs. Larger slopes indicated a sharper and less variable transition between response categories. Importantly, psychometric slope was not assumed to provide a process-pure measure of sensory temporal precision, as its magnitude may reflect variability arising from both sensory encoding and decisional processes. Response asymptotes were therefore modelled separately to account for occasional stimulus-independent responses, such as attentional lapses, accidental responses, or motor errors, that could otherwise influence estimates of the psychometric function^60,82^. In models containing a symmetric lapse parameter, the total participant-specific lapse probability was divided equally between the lower and upper asymptotes. Thus, half of the estimated lapse probability was assigned to each asymptote. Candidate models with asymmetric response errors instead estimated separate participant-specific lower and upper asymptotic errors. Furthermore, to assess the influence of recent perceptual experience on current temporal judgements, trials were classified according to the perceptual outcome reported on the immediately preceding trial (t−1 integration and t−1 segregation). The modelling framework therefore estimated PSE and psychometric slope while simultaneously accounting for both perceptual-history effects and stimulus-independent responses. History-neutral PSE and slope parameters were estimated together with their corresponding history-dependent values following t−1 integration and t−1 segregation. Perceptual-history effects were quantified as the difference between the t−1 integration and t−1 segregation conditions for both PSE and slope. For PSE, larger positive differences reflected stronger attractive serial dependence, values around zero indicated little systematic influence of perceptual history, and negative differences indicated a repulsive history effect.

#### Visual temporal performance: Hierarchical model structure and candidate models

Psychometric parameters were estimated hierarchically within a Bayesian framework, allowing partial pooling across participants while retaining participant-specific variation. Participant-level estimates were therefore informed by the corresponding group-level distributions while preserving meaningful individual differences^83^. Participant-specific deviations were implemented using non-centred parameterizations to improve sampling efficiency and facilitate stable estimation. Weakly informative priors were specified for all group-level parameters and between-participant variability terms. The history-neutral PSE was assigned a Normal prior centred at 75 ms with a standard deviation of 35 ms, whereas between-participant variability was modelled using a Half-Normal prior with a scale of 25 ms. The history-dependent PSE effect was assigned a zero-centred Normal prior with a standard deviation of 25 ms, with between-participant variability assigned a Half-Normal prior with a scale of 20 ms. Psychometric slope was estimated on the logarithmic scale, with the history-neutral group-level log slope assigned a Normal prior centred at log(0.07) with a standard deviation of 0.70 and its between-participant variability a Half-Normal prior with a scale of 0.50. The history effect on log slope was assigned a zero-centred Normal prior with a standard deviation of 0.35, with between-participant variability assigned a Half-Normal prior with a scale of 0.35, reflecting no a priori directional assumption regarding perceptual-history effects. For models containing a symmetric lapse parameter, lapse probability was estimated on the logit scale using a Normal prior centred at logit(0.06) with a standard deviation of 1.00, with participant-level variability assigned a Half-Normal prior with a scale of 0.75. Candidate models containing asymmetric response errors used corresponding priors for the lower and upper asymptotic-error parameters. Prior predictive checks were performed for the final model to assess whether the specified priors generated plausible psychometric response patterns before conditioning on the observed response counts. We generated 2,000 samples from the prior predictive distribution, computing the prior-predictive median and 95% highest-density interval. The resulting prior-predictive distributions spanned a broad range of plausible psychometric functions across the full response-probability range while retaining the expected monotonic transition from integration to segregation with increasing ISI. All pooled observed response proportions fell within the corresponding 95% prior-predictive intervals, indicating that the priors were sufficiently broad to encompass the observed behavioural patterns without imposing overly restrictive expectations on the psychometric function.

We fitted 12 candidate hierarchical psychometric models forming a complete 2 × 2 × 3 factorial model space. Importantly, the model space included baseline models in which neither PSE nor psychometric slope varied as a function of perceptual history, allowing us to test whether incorporating perceptual-history effects improved the characterization of behavioural performance rather than assuming their presence a priori. Models then systematically tested all combinations in which perceptual history affected the PSE only, the psychometric slope only, or both PSE and slope. Each of these four psychometric structures was evaluated under three alternative specifications of the response asymptotes: fixed at zero and one, governed by a symmetric participant-specific lapse parameter, or estimated using separate participant-specific lower and upper asymptotic errors. This allowed us to formally test whether accounting for occasional stimulus-independent responses improved the characterization of behavioural performance relative to a standard psychometric function with fixed asymptotes, and whether such responses were adequately captured by a single symmetric lapse parameter or required separate lower and upper asymptotic-error terms. Accordingly, the 12 candidate models were:

- M1: shared PSE and shared slope, with fixed response asymptotes;
- M2: history-dependent PSE and shared slope, with fixed response asymptotes;
- M3: shared PSE and history-dependent slope, with fixed response asymptotes;
- M4: history-dependent PSE and history-dependent slope, with fixed response asymptotes;
- M5: shared PSE and shared slope, with a symmetric participant-specific lapse parameter;
- M6: history-dependent PSE and shared slope, with a symmetric participant-specific lapse parameter;
- M7: shared PSE and history-dependent slope, with a symmetric participant-specific lapse parameter;
- M8: history-dependent PSE and history-dependent slope, with a symmetric participant-specific lapse parameter;
- M9: shared PSE and shared slope, with separate participant-specific lower and upper asymptotic errors;
- M10: history-dependent PSE and shared slope, with separate participant-specific lower and upper asymptotic errors;
- M11: shared PSE and history-dependent slope, with separate participant-specific lower and upper asymptotic errors;
- M12: history-dependent PSE and history-dependent slope, with separate participant-specific lower and upper asymptotic errors.

Thus, the factorial model space allowed us to evaluate perceptual-history effects on PSE and psychometric slope independently, while simultaneously testing alternative assumptions about stimulus-independent response errors. Whenever a perceptual-history effect was included, both a group-level effect and participant-specific hierarchical variation were estimated.

#### Visual temporal performance: Model estimation, diagnostic and comparison

Each model was estimated using four Markov chain Monte Carlo chains, with 3,000 tuning iterations followed by 3,000 retained posterior draws per chain. The target acceptance probability was set to 0.97 and the maximum tree depth to 15. Candidate models were compared using Pareto-smoothed importance-sampling leave-one-out cross-validation. Exact leave-one-out refitting was performed for observations with Pareto *k* > .70. Before entering model comparison, each model was required to pass convergence, effective-sample-size, divergent-transition, tree-depth, and Pareto-k diagnostics. When multiple models were predictively indistinguishable according to a prespecified one-standard-error criterion, the model with the lower complexity was selected. The asymmetric-asymptote model containing perceptual-history effects on both PSE and slope produced the highest estimated out-of-sample predictive accuracy. However, the corresponding model containing a symmetric participant-specific lapse parameter was within one standard error of this more complex model (ΔELPD_LOO_ =1.83, SE = 5.66) and was therefore selected as the more parsimonious model. Full model-comparison results and sampling diagnostics for all 12 candidate models are reported in Supplementary Table S1. The selected model consequently included participant-specific effects of perceptual history on both PSE and psychometric slope, together with a participant-specific symmetric lapse parameter. All convergence and sampling diagnostics for the selected model were satisfactory. The maximum R-hat was 1.00, the minimum bulk and tail effective sample sizes were 1,354 and 2,454, respectively, no divergent transitions occurred, and no iterations reached the maximum tree depth. Visual inspection of the chain-specific posterior distributions and MCMC trace plots further supported optimal convergence and sampling stability, with strong overlap across chains, well-mixed traces throughout the retained posterior draws (Fig. S2). Posterior predictive checks were then used to assess whether the selected model adequately reproduced the observed behavioural data. Observed response counts fell within the corresponding 95% posterior-predictive interval. At the group level, posterior-predicted psychometric functions closely reproduced the observed probabilities of segregation across ISIs for both t-1 integration and t-1 segregation trials (Fig. S3), indicating that the selected model provided an accurate characterization of the observed history-dependent psychometric responses.

#### Subjective confidence: Hierarchical Bayesian modelling

Confidence ratings were analyzed using a trial-level hierarchical Bayesian ordered-logistic model, preserving the original four-level confidence scale (1–4) as an ordinal outcome. To determine whether individual differences in reported confidence reflected general differences in confidence-scale use and the extent to which confidence tracked perceptual evidence, we modelled confidence at the trial level while separating participant-specific baseline confidence from evidence-dependent modulation. To quantify perceptual-evidence strength, we derived an individualized measure of perceptual evidence from the selected hierarchical psychometric model described above. For each participant, history-neutral signed perceptual evidence was calculated from the posterior median participant-specific PSE and psychometric slope. Negative values therefore represented evidence favouring perceptual integration (individual integration window), positive values represented evidence favouring segregation (individual segregation window), and zero corresponded to the participant-specific history-neutral PSE. Signed evidence was scaled by its pooled within-participant standard deviation without subtracting participant-specific means, thereby preserving zero at the participant-specific PSE. Absolute evidence was defined as the magnitude of signed evidence, centred within participant, and divided by its pooled within-participant standard deviation. Absolute evidence therefore quantified the strength of current perceptual evidence irrespective of whether it favoured integration or segregation. Because absolute evidence was centred within participant, its effect quantified whether confidence changed with trial-to-trial variations in evidence strength within individuals, independently of stable between-participant differences in average evidence. To test whether recent perceptual experience modulated confidence, a perceptual history-congruence predictor was constructed from signed perceptual evidence and the direction of the preceding percept, and multiplied by signed evidence. Positive history-congruence values therefore indicated that current perceptual evidence favoured the same perceptual interpretation reported on the preceding trial, whereas negative values indicated that current evidence favoured the opposite interpretation. The model simultaneously included population-level effects of absolute evidence, signed evidence, current perceptual response, preceding perceptual response, history congruence, and trial position. Trial position was centred within participant and standardized by its pooled within-participant standard deviation to account for systematic changes in confidence over the course of the experiment. Thus, the absolute-evidence effect tested whether confidence systematically tracked the strength of individualized perceptual evidence, whereas the history-congruence effect tested whether confidence was selectively enhanced when current evidence supported the perceptual interpretation experienced on the preceding trial.

#### Subjective confidence: Hierarchical model structure and priors

Confidence parameters were estimated hierarchically, allowing partial pooling across participants while retaining participant-specific variation. Correlated participant-specific random effects were included for the intercept, absolute-evidence slope, and history-congruence slope. The participant-specific intercept captured individual differences in the general tendency to use higher or lower portions of the confidence scale, conditional on the remaining model predictors. In contrast, the participant-specific absolute-evidence slope captured individual differences in the extent to which confidence tracked within-participant changes in perceptual evidence strength. The participant-specific history-congruence slope captured individual differences in the extent to which confidence was modulated by consistency between current perceptual evidence and the preceding percept. Estimating these components simultaneously therefore allowed general confidence tendency to be distinguished from evidence-dependent and history-dependent confidence modulation. The three participant-specific effects were estimated jointly, allowing their between-participant variation to be correlated. Weakly informative priors were specified for all population-level and participant-level parameters. The population-level absolute-evidence effect was assigned a zero-centred Normal prior with a standard deviation of 1.00. Signed evidence, current perceptual response, preceding perceptual response, and history congruence were each assigned zero-centred Normal priors with a standard deviation of 0.75, whereas trial position was assigned a zero-centred Normal prior with a standard deviation of 0.50. Participant-level standard deviations for the intercept, absolute-evidence slope, and history-congruence slope were assigned Half-Normal priors with a scale of 1.00. The three ordered-logistic cutpoints separating the four confidence categories were assigned ordered Normal priors centred at −1.5, 0, and 1.5, respectively, with a standard deviation of 2.5. Prior predictive checks were performed from the joint prior predictive distribution to determine whether the specified priors generated plausible confidence-response distributions before conditioning on the observed data. The observed proportions of all four confidence categories fell within their corresponding 95% prior-predictive intervals, indicating that the priors were sufficiently broad to encompass the observed confidence distribution without imposing overly restrictive expectations.

#### Subjective confidence: Model estimation and posterior inference

Four Markov chain Monte Carlo chains were estimated, with 3,000 tuning iterations followed by 3,000 retained posterior draws per chain. All convergence and sampling diagnostics were satisfactory. The maximum R-hat was 1.009, the minimum bulk and tail effective sample sizes were 746 and 1,599, respectively, no divergent transitions occurred, and no iterations reached the maximum tree depth. Visual inspection of chain-specific posterior distributions and MCMC trace plots further supported optimal convergence and sampling stability, with well-mixed and stationary chains across the retained posterior draws (Fig. S4). Posterior predictive checks were performed using 400 posterior-predictive replications, based on 100 posterior draws sampled from each of the four chains. Model adequacy was evaluated at multiple levels, including reproduction of the overall distribution of the four ordinal confidence categories, the relationship between confidence and absolute perceptual-evidence strength, and the history-dependent relationship between confidence and signed perceptual evidence. Posterior-predicted confidence patterns closely reproduced the principal features of the observed data across these dimensions (Fig. S5), supporting the adequacy of the hierarchical model for characterizing variations in confidence. The original ordinal confidence ratings were used for all model estimation and statistical inference. Within-participant standardized confidence values were calculated only for visualization of the history-dependent confidence effect and did not enter the ordered-logistic model.

### EEG analysis

Resting-state EEG was recorded in two conditions, eyes closed (EC) and eyes open (EO), each lasting 3 min. Signals were acquired from 64 active scalp electrodes using BrainVision Recorder (Brain Products), sampled at 1000 Hz, and referenced online to FCz. Impedances were kept below 10 kΩ. Participants were instructed to relax, minimize movements, and fixate (EO) or keep their eyes closed (EC). Preprocessing was performed in MATLAB. For each participant, noisy channels were identified by visual inspection and interpolated using spherical splines. Oculomotor artifacts were removed by independent component analysis (ICA). The reference was kept at FCz for all subsequent analyses. We intentionally avoided re-referencing to the average reference because broadband noise or a single noisy channel would otherwise be spread over all channels and distort the estimation of the aperiodic (1/f) component^27^. For spectral analysis, continuous resting-state data were segmented into 1-s epochs. Remaining segments were submitted to Welch’s method, yielding a power spectral density (PSD) estimate. To separate oscillatory from aperiodic activity, we applied the “fitting oscillations and one-over-f” (FOOOF) algorithm^32^ to the PSD of each participant, condition (EC, EO), and electrode. FOOOF models the log-power spectrum as the sum of (i) an aperiodic component following a 1/f function (offset and exponent) and (ii) a set of Gaussian peaks capturing periodic oscillations. Default settings were used (peak width limits: 0.5– 12 Hz; minimum peak height: 0; maximum number of peaks: Inf; peak threshold: 2). The aperiodic exponent (slope) from this fit was taken as our index of aperiodic neural activity.

To quantify oscillatory activity independently of the aperiodic background, we derived the aperiodic-corrected spectrum by subtracting the fitted aperiodic component from the log-transformed PSD. Individual alpha frequency (IAF) was defined as the frequency exhibiting maximum aperiodic-corrected spectral power within the alpha range (8–13 Hz). As frequency-based control analyses, analogous dominant frequency estimates were extracted as the frequency corresponding to the maximum of the aperiodic-corrected spectrum within the delta (1–4 Hz), theta (4–7 Hz), low-beta (15–30 Hz), high-beta (30–45 Hz), and gamma (45–70 Hz) ranges. In addition, to examine whether behavioural relationships were associated with oscillatory power rather than dominant frequency, we quantified aperiodic-corrected periodic power as the mean corrected spectral power within each of the same canonical frequency bands described above. In parallel, periodic peaks independently identified by FOOOF were characterized within each frequency band according to their centre frequency. Prevalence was calculated across all participant–electrode spectra, with one spectrum estimated separately for each participant and scalp electrode. Alpha peaks (8–13 Hz) showed particularly high reliability, being identified in 99.4% and 98.1% of spectra in EC and EO, respectively. Peak prevalence was more heterogeneous outside the alpha range: delta peaks (1–4 Hz) were identified in 10.6% and 16.5% of spectra in EC and EO, respectively, theta peaks (4–7 Hz) in 46.8% and 54.4%, beta peaks (15–45 Hz) in 60.45% and 60.2%, and gamma peaks (45–70 Hz) in 76.1% and 78.5%. As an additional validation of the aperiodic-corrected IAF used as our primary oscillatory frequency measure, corrected-spectrum IAF estimates were compared with the centre frequency of alpha peaks independently identified by FOOOF. Agreement was very high across electrodes in both EC (r = 0.975) and EO (r = 0.976).

### Behavioral Statistical Analysis

For visual temporal performance, statistical inference on temporal integration thresholds, psychometric slopes, and perceptual-history effects was based on the posterior distributions of the hierarchical Bayesian psychometric model. Group-level effects were summarized using posterior estimates and 95% highest-density intervals (HDIs), with directional evidence additionally quantified as the posterior probability that the corresponding effect was greater or smaller than zero. Perceptual-history effects were quantified as the difference between trials preceded by integration and segregation. For PSE, this contrast was defined as ΔPSE = PSE t−1integration − PSE t−1segregation, with larger positive values indicating stronger attractive serial dependence. History-dependent differences in psychometric slope were estimated on the log-slope scale as the corresponding contrast between t−1 integration and t−1 segregation conditions. For subsequent individual-differences analyses, participant-level posterior means from the hierarchical model were used as summary estimates and used in subsequent correlational analyses with EEG-derived parameters.

For confidence analyses, mean confidence was first related to participant-specific PSE and psychometric slope estimates using Pearson correlations. For the model-free perceptual-history analysis, trial-wise confidence ratings were sorted according to ISI and preceding perceptual outcome (t−1 integration versus t−1 segregation). Confidence scores were z-scored within participants to preserve within-participant relative variability while removing stable between-participant differences in overall confidence level. Differences in confidence between t−1 integration and t−1 segregation trials were assessed using permutation-based paired-sample t-tests conducted separately for each ISI. In each permutation (N = 10,000), participant-wise differences were randomly sign-flipped and a new t-statistic was computed to generate an empirical null distribution. Two-tailed empirical p-values were defined as the proportion of permuted t-values with an absolute magnitude greater than or equal to the observed t-statistic. To correct for multiple comparisons across ISIs, p-values were adjusted using the Benjamini–Hochberg false discovery rate procedure (q = 0.05). Effect sizes were calculated as Cohen’s d for paired samples.

Statistical inference for the hierarchical Bayesian confidence model was based on the posterior distributions of the population-level coefficients and participant-level random effects. Effects were summarized using posterior estimates and 95% HDIs, with directional evidence additionally quantified as the posterior probability that the corresponding coefficient was greater or smaller than zero. For ordered-logistic population-level coefficients, odds ratios were obtained by exponentiating the corresponding posterior draws. Between-participant variation in general confidence tendency and in the strength of confidence–evidence and history-congruence coupling was quantified from the posterior distributions of the corresponding participant-level random-effect standard deviations.

### Resting-State EEG Statistical Analysis

To characterize the relationship between eyes-closed and eyes-open resting-state spectral activity, we investigated state-dependent differences for our primary measures of interest: IAF (aperiodic-corrected) and aperiodic exponent. EC–EO differences were assessed at each electrode using paired, two-sided cluster-based permutation tests (10,000 permutations)^84^. For each participant and electrode, the difference was defined as EC − EO. Paired *t*-statistics were calculated across participants, and spatial clusters were formed using a two-sided channel-level threshold of p < 0.05 and triangulation-based spatial neighbourhoods. Cluster mass was defined as the summed t-statistic within each spatial cluster, and cluster-level significance was assessed against the maximum cluster-mass distribution obtained from 10,000 Monte Carlo permutations. To assess whether individual differences in resting-state spectral activity across EC and EO, we additionally calculated Pearson correlations between EC and EO values across participants at each electrode. Statistical inference was again performed using two-sided spatial cluster-based permutation testing (10,000 permutations). All EEG analyses were implemented in MATLAB using custom scripts in combination with the FieldTrip Toolbox^85^ and the FOOOF toolbox^32^.

### Behavior – Resting-State EEG: Statistical Analysis

#### Alpha Frequency and Aperiodic Predictors of Temporal Integration

For each resting-state condition (eyes closed, EC; eyes open, EO), EEG–behaviour relationships were assessed at the scalp level using nonparametric cluster-based permutation statistics^84^. First, participant-level posterior mean estimates of the psychometric threshold (PSE) and psychometric slope derived from the hierarchical Bayesian psychometric model were correlated with IAF and the aperiodic exponent at each electrode. Associations were tested using Pearson correlations, and clusters were formed using a two-sided channel-level threshold of p < 0.05 and triangulation-based spatial neighbourhoods. Cluster mass was defined as the summed t-statistic within each spatial cluster, and cluster-level significance was assessed against the maximum cluster-mass distribution obtained from 10,000 Monte Carlo permutations. To additionally control for the number of scalp-level correlational analyses performed, the minimum cluster-corrected p-value from each analysis (PSE and psychometric slope × EC and EO) was subsequently entered into a Benjamini–Hochberg false-discovery-rate (FDR) correction^86^, with statistical significance defined as an FDR-adjusted q < .05. As an additional control, participant-specific lapse-rate estimates from the hierarchical psychometric model were correlated with resting-state IAF and aperiodic exponent separately for EC and EO using the same scalp-level cluster-permutation framework applied to the primary EEG–behavior analyses. All correlation analyses were performed separately for eyes-closed (EC) and eyes-open (EO) resting-state measures, with the corresponding topographic maps and scatterplots likewise presented separately for each resting-state condition. Additionally, to assess whether the association between resting IAF and temporal integration (PSE) could be explained by frequency correspondence between the endogenous alpha rhythm and the rhythmic structure of our sustained stimulus sequence^65^, we performed an additional hierarchical Bayesian analysis explicitly accounting for both resting IAF and stimulus frequency–IAF distance. Analyses were performed separately using eyes-closed (EC) and eyes-open (EO) resting-state IAF. For each participant and resting-state condition, IAF was defined as the mean aperiodic-corrected alpha peak frequency across a predefined posterior ROI (O1, Oz, O2, PO3, POz, PO4, PO7, PO8) and was standardized across participants within condition. In the baseline model (IAF-only model), standardized IAF was entered as a participant-level predictor of PSE, such that the corresponding coefficient, β_IAF_, quantified the change in PSE, in milliseconds, associated with a 1-SD increase in resting IAF. To quantify stimulus frequency–IAF correspondence, the stimulus rate associated with each non-zero ISI condition was derived from the corresponding stimulus duration. We focused on stimulus frequencies falling within the same 8–13 Hz range used to define IAF, corresponding to 12, 9.6, and 8 Hz (83.33, 104.17, and 125 ms stimulus durations, respectively). For each participant, stimulus frequency–IAF distance was initially defined as the absolute difference between each of these frequencies and the participant’s resting IAF. These distance values were then two-way centered across participants and stimulus frequencies, removing both each participant’s average distance and each stimulus frequency’s average distance while retaining the participant-by-frequency correspondence component. All behavioral observations remained in the likelihood, whereas timing conditions outside the predefined alpha range, including the non-rhythmic 0-frame fused control, were assigned a distance contribution of zero. An extended model (IAF + distance model) added a single population-level distance coefficient to the baseline model, allowing the local PSE to vary as a function of the centered stimulus frequency–IAF distance. Thus, β_distance_ quantified the change in PSE associated with a 1-Hz increase in relative stimulus frequency–IAF distance. The IAF coefficient was assigned a zero-centered Normal prior with SD = 15 ms per 1-SD IAF, and the distance coefficient a zero-centered Normal prior with SD = 10 ms/Hz. Prior-predictive checks were performed before posterior fitting and confirmed that these priors generated broad, plausible distributions of PSEs and behavioral responses. Each model was estimated with four MCMC chains, 3,000 tuning iterations and 3,000 retained draws per chain. Models were assessed using convergence and sampling diagnostics together with posterior predictive checks. All models showed satisfactory convergence and sampling performance (R^ = 1.00, minimum bulk effective sample size > 1,900, no divergences, and no maximum tree-depth transitions). Posterior-predictive checks further indicated good model fit (Fig. S6). The IAF-only and IAF plus distance models were additionally compared using Pareto-smoothed importance-sampling leave-one-out cross-validation. Observations with Pareto k > .70 were evaluated using exact leave-one-out refitting. The primary inferential measures were the posterior distributions of β_IAF_ and β_distance_, together with the change in the IAF coefficient after inclusion of stimulus frequency–IAF distance.

#### Alpha Frequency and Aperiodic Predictors of Serial Dependence and Confidence

Next, to assess whether EEG parameters predicted the strength of serial dependence, we further correlated EEG measures with ΔPSE (t–1 integration -t–1 segregation). ΔPSE was correlated separately with IAF and the aperiodic exponent in EC and EO using the same two-sided cluster-based Pearson-correlation framework described above. Given that both IAF and aperiodic exponent predicted serial dependence strength (ΔPSE) in the EO resting-state condition, we conducted reciprocal mediation analyses using the lavaan package (version 0.7–2) in RStudio. All variables were z-standardized prior to analysis to facilitate interpretation of path coefficients. Structural equation models were estimated using maximum likelihood estimation with Full

Information Maximum Likelihood (FIML). Indirect effects were assessed using 10,000 bootstrap resamples, with bias-corrected 95% confidence intervals (CIs), and path coefficients are reported as standardized regression weights (β). IAF and aperiodic exponent were averaged across the electrodes belonging to significant EO correlation clusters with ΔPSE. Effects were considered reliable when the 95% bootstrap CI of the indirect effect did not include zero. Inference focused on the direct, indirect, and total effects, together with the proportion of variance explained in ΔPSE (R²).

For subjective confidence, we first tested whether resting-state EEG predicted participants’ overall mean confidence ratings. Mean confidence was correlated separately with IAF and the aperiodic exponent in EC and EO using Pearson’s correlations with cluster-based nonparametric correction (10,000 permutations) using the same framework described above. We next examined whether EEG predicted the two participant-specific confidence dimensions estimated by the hierarchical Bayesian ordered-logistic model: general confidence tendency, corresponding to the participant-specific random intercept, and confidence–evidence coupling, corresponding to the participant-specific absolute-evidence slope. Because these parameters may share between-participant variance, their neural associations were assessed using reciprocal scalp-level partial correlations. For the general-confidence-tendency analysis, the association between EEG and general confidence tendency was estimated while controlling for confidence–evidence coupling. Conversely, for the confidence–evidence-coupling analysis, general confidence tendency was treated as the covariate. Partial correlations were implemented using Frisch–Waugh–Lovell residualization, whereby the nuisance confidence component was removed from both the target confidence parameter and the EEG measure at each electrode before computing their correlation. Statistical significance for these partial-correlation maps was assessed using a Freedman–Lane permutation procedure. Reduced-model residuals were permuted across participants using the same permutation at all electrodes, preserving the spatial dependence of the EEG data. Two-sided clusters were formed at p < 0.05 using the same triangulation-based electrode neighbourhood structure. Cluster mass was quantified as the sum of the absolute t-statistics within spatially connected, same-sign clusters, and the maximum cluster mass across positive and negative clusters was retained for each of 10,000 permutations to construct the null distribution.

#### Control Analyses: Frequency Band Specificity

To assess whether the observed EEG–behaviour associations were specific to the spectral features of primary interest, we performed complementary frequency-and power-based control analyses using the same scalp-level cluster-permutation framework described above. For the frequency-based controls, peak frequency estimates (aperiodic-corrected) were defined as the frequency corresponding to the maximum corrected spectral power within the delta (1–4 Hz), theta (4–7 Hz), low beta (15–30 Hz), high beta (30-45), and gamma (45–70 Hz) ranges. These estimates were derived independently of formal FOOOF peak detection and therefore provided an analogous frequency-based measure across all participants and electrodes. For the power-based controls, aperiodic-corrected oscillatory power was for each frequency band, thereby assessing band-limited periodic activity after removal of the fitted aperiodic background. For PSE and psychometric slope, the five non-alpha peak-frequency measures were tested separately in EC and EO, yielding 20 scalp-level analyses (5 frequency bands × 2 behavioural measures × 2 resting-state conditions). Aperiodic-corrected power was tested analogously across six frequency bands, yielding 24 analyses (6 bands × 2 behavioural measures × 2 conditions). Analogous spectral-specificity controls were performed for Confidence analyses. These control analyses were restricted to Mean

Confidence and were not extended to General Confidence Tendency or Confidence–Evidence Coupling, which were examined separately using the focused reciprocal partial-correlation analyses described above.

## Acknowledgements

This work was supported by the NYUAD Center for Brain and Health, funded by Tamkeen under NYU Abu Dhabi Research Institute grant CG012.

## Author contributions

Conceptualization, G.M., M.D., D.M.; Methodology G.M., M.D., D.M.; Formal Analysis, G.M.; Software, G.M.; Visualization, G.M.; Funding Acquisition, D.M.; Writing -Original Draft Preparation, G.M. D.M.; Writing – Review & Editing, G.M., M.D., D.M.; Supervision, D.M.

## Data availability

All data and code are openly available on the Open Science Framework (OSF) at https://osf.io/4wdmc/.

## Notes

### Competing Interest Statement

The authors have declared no competing interest.

### Summary of Updates

This revised manuscript incorporates all modifications made in response to the journal's peer-review comments.

