## Supplementary Information for "Alpha oscillations and aperiodic neural dynamics jointly predict visual temporal resolution, confidence, and dependence on prior experience"

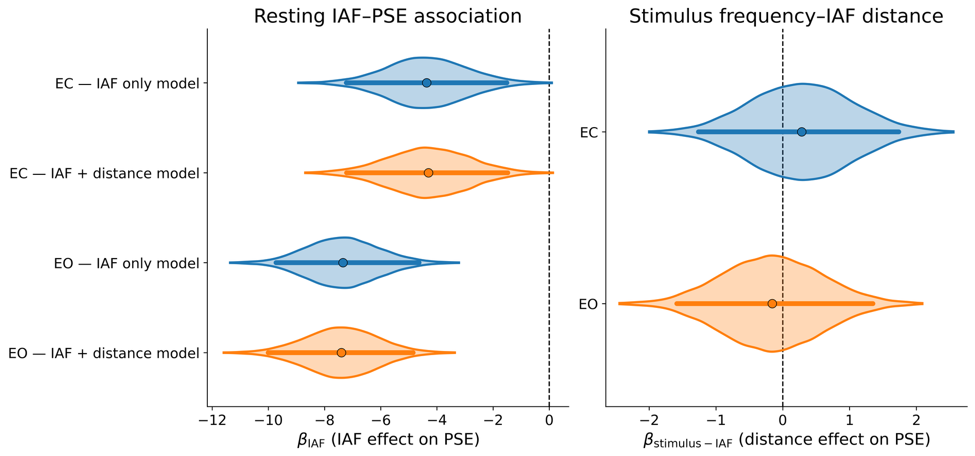
 **Figure S1. Resting IAF–PSE association after accounting for stimulus frequency–IAF distance.** Left panel, posterior distributions of the resting IAF coefficient (βIAF) from the IAF-only model and from the extended IAF plus distance model, shown separately for eyes-closed (EC) and eyes-open (EO) resting-state IAF. βIAF quantifies the change in temporal-integration PSE, in milliseconds, associated with a 1-SD increase in resting posterior IAF; negative values therefore indicate lower PSEs, and hence narrower temporal integration windows, at faster IAF. Right, posterior distributions of the stimulus frequency–IAF distance coefficient (βstimulus–IAF) from the extended model. Stimulus frequency–IAF distance was defined for the three stimulus-duration conditions whose alternation rates fell within the alpha range (8–13 Hz): 83.33 ms (12 Hz), 104.17 ms (9.6 Hz), and 125 ms (8 Hz), and was two-way centered across participants and stimulus frequencies before model inclusion. βstimulus–IAF therefore quantifies the change in PSE associated with greater relative stimulus frequency–IAF distance. Dots indicate posterior means and thick horizontal lines indicate 95% highest-density intervals (HDIs); dashed vertical lines denote zero effect. Shaded densities show the central 99.5% of posterior samples for visualization, whereas posterior means and HDIs were calculated from the complete posterior distributions. The IAF–PSE association remained credibly negative and essentially unchanged after inclusion of stimulus frequency–IAF distance, whereas the distance-effect HDIs spanned zero for both EC and EO.

**Hierarchical Bayesian Psychometric Model Diagnostic**

| **Model** | **Model structure** | **ELPD_LOO_** | **p_LOO_** | **max R-hat** | **min ESS bulk** | **min ESS tail** | **min BFMI** |
| --- | --- | --- | --- | --- | --- | --- | --- |
| 1 | Shared PSE + shared slope + fixed asymptotes | -2062.16 | 311.86 | 1.00 | 636 | 1398 | .641 |
| 2 | Shared PSE + history-dependent slope + fixed asymptotes | -2071.49 | 382.30 | 1.00 | 549 | 1230 | .626 |
| 3 | History-dependent PSE + shared slope + fixed asymptotes | -1671.97 | 259.80 | 1.01 | 809 | 1859 | .622 |
| 4 | History-dependent PSE + history-dependent slope + fixed asymptotes | -1665.47 | 289.13 | 1.00 | 753 | 1699 | .617 |
| 5 | Shared PSE + shared slope + symmetric lapse | -2040.28 | 342.03 | 1.00 | 1163 | 2085 | .686 |
| 6 | Shared PSE + history-dependent slope + symmetric lapse | -2052.09 | 385.69 | 1.00 | 1472 | 2010 | .664 |
| 7 | History-dependent PSE + shared slope + symmetric lapse | -1631.74 | 259.96 | 1.00 | 1421 | 2936 | .650 |
| **8** | **History-dependent PSE + history-dependent slope + symmetric lapse (selected)** | **-1623.61** | **279.91** | **1.00** | **1354** | **2454** | **.624** |
| 9 | Shared PSE + shared slope + asymmetric asymptotes | -2039.88 | 362.88 | 1.00 | 1004 | 2166 | .718 |
| 10 | Shared PSE + history-dependent slope + asymmetric asymptotes | -2038.37 | 384.25 | 1.00 | 890 | 1761 | .653 |
| 11 | History-dependent PSE + shared slope + asymmetric asymptotes | -1633.21 | 264.89 | 1.00 | 1101 | 2718 | .662 |
| 12 | History-dependent PSE + history-dependent slope + asymmetric asymptotes | -1621.78 | 280.61 | 1.00 | 1185 | 1932 | .674 |

**Table S1. Hierarchical Bayesian psychometric model comparison and sampling diagnostics.** Twelve candidate hierarchical psychometric models were compared within a complete 2 × 2 × 3 factorial model space, systematically varying whether the Point of Subjective Equality (PSE) and psychometric slope were shared across perceptual-history conditions, and whether response asymptotes were fixed at zero and one, governed by a symmetric participant-specific lapse parameter, or estimated using separate participant-specific lower and upper asymptotic errors. Models are presented in the same order as the prespecified candidate model space, whereas Rank indicates their ordering according to expected out-of-sample predictive performance. ELPD_LOO_ denotes the expected log pointwise predictive density estimated using Pareto-smoothed importance-sampling leave-one-out cross-validation, with higher values indicating better predictive performance; p_LOO_ denotes the effective number of parameters. Sampling diagnostics include the maximum potential scale-reduction factor (R-hat), minimum bulk and tail effective sample sizes (ESS), and minimum Bayesian fraction of missing information (BFMI). All candidate models showed satisfactory sampling diagnostics, with no divergent transitions and no iterations reaching the maximum tree depth. Although the model with history-dependent PSE and slope and asymmetric response asymptotes achieved the highest ELPD_LOO_, the corresponding model with a symmetric participant-specific lapse parameter was predictively indistinguishable according to the prespecified one-standard-error criterion (ΔELPD_LOO_ = 1.83, SE = 5.66) and was therefore selected as the more parsimonious model.

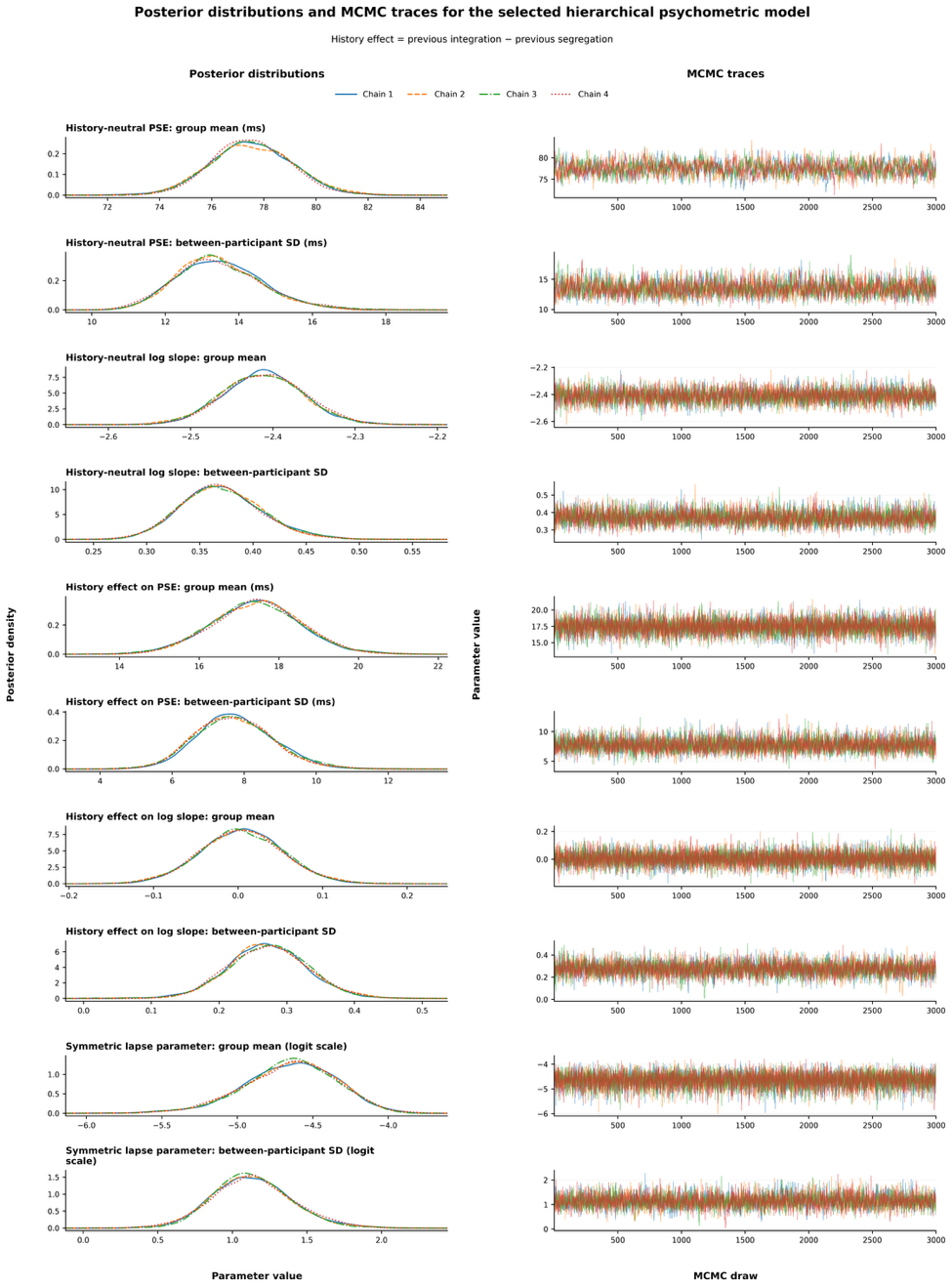

**Supplementary Figure S2. Posterior distributions and MCMC traces for the selected hierarchical psychometric model.** Posterior distributions and Markov chain Monte Carlo traces are shown for the principal group-level parameters of the selected hierarchical psychometric model. The left column shows chain-specific posterior density estimates, and the right column shows the corresponding sampling traces across 3,000 post-warm-up draws for each of four chains. Parameters include the group mean and between-participant standard deviation of the history-neutral point of subjective equality (PSE), the history-neutral log slope, the history effect on PSE, the history effect on log slope, and the symmetric lapse parameter. History effects are defined as the parameter value following a previous integration report minus the corresponding value following a previous segregation report. The strong overlap of the chain-specific posterior distributions and the stationary, well-mixed traces indicate satisfactory convergence and sampling stability.

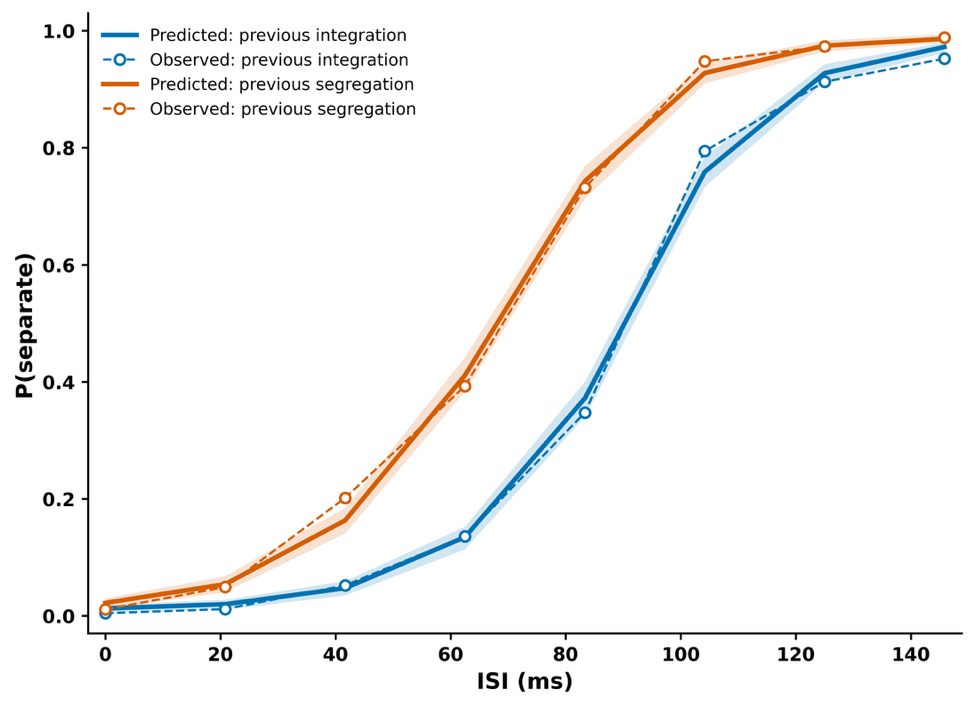
 **Figure S3. Posterior predictive check for the selected hierarchical psychometric model.** Observed and posterior-predicted probabilities of reporting the current stimulus as separate are shown as a function of the current interstimulus interval (ISI), separately according to the percept reported on the preceding trial. Blue curves indicate trials preceded by an integrated percept, whereas orange curves indicate trials preceded by a segregated percept. Solid lines show posterior-predictive means, shaded regions show 95% highest-density intervals, and dashed lines with open circles show the observed response proportions. The close correspondence between observed and predicted values indicates that the selected model adequately reproduces the history-dependent psychometric functions across ISIs.

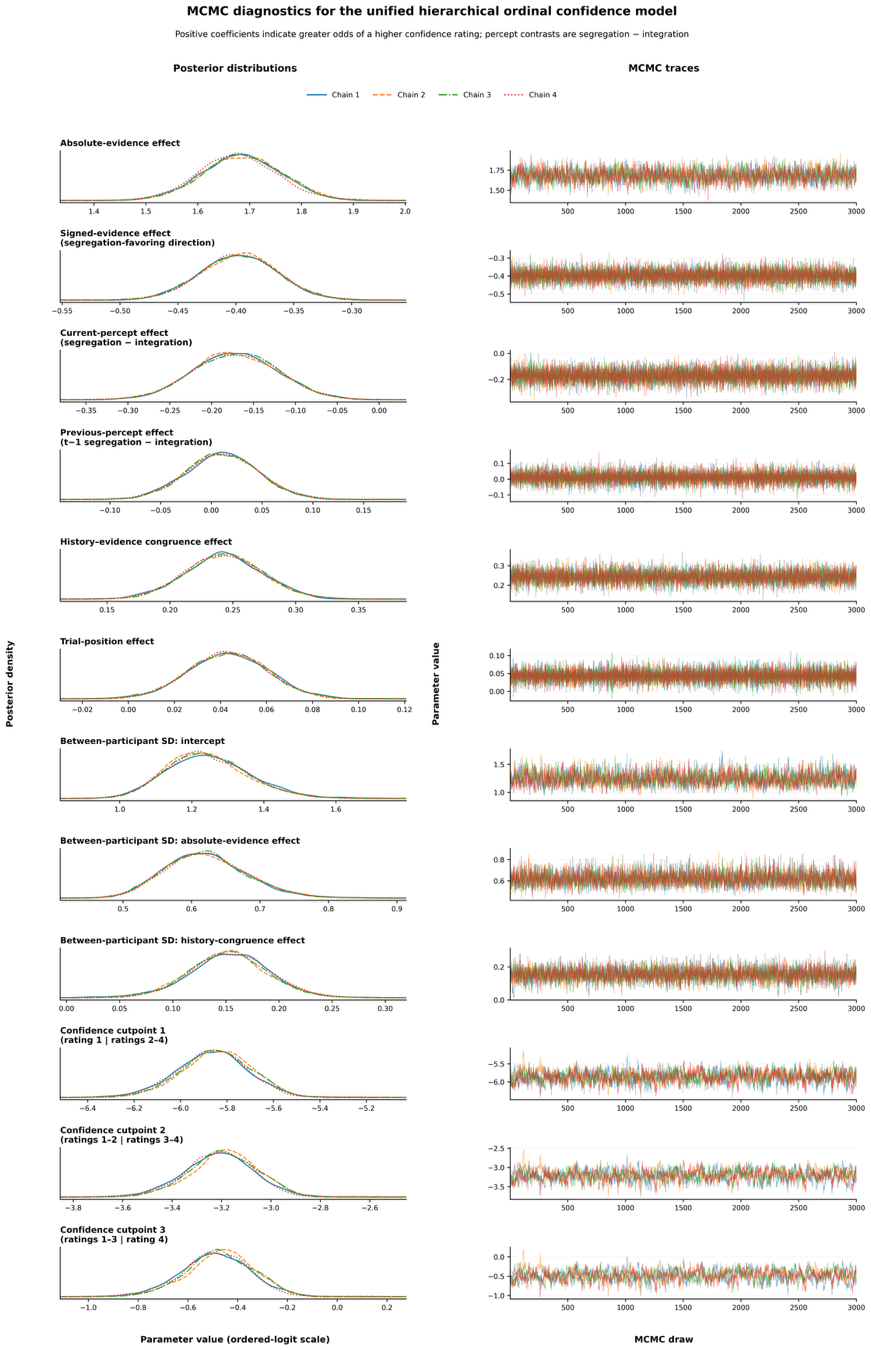

**Figure S4. Posterior distributions and MCMC sampling traces for the hierarchical ordinal confidence model.** Chain-specific posterior distributions and sampling traces are shown for the population-level effects, between-participant standard deviations, and ordered-logistic confidence cutpoints of the model. The left column shows posterior density estimates, and the right column shows the corresponding traces across 3,000 draws for each of four chains. Positive population-level coefficients indicate greater odds of reporting a higher confidence category. Current- and previous-percept effects are coded as segregation minus integration, while positive history–evidence congruence values indicate that the current sensory evidence supports the perceptual interpretation reported on the preceding trial. Overlapping chain-specific posterior distributions and stationary traces are consistent with satisfactory convergence and sampling stability.

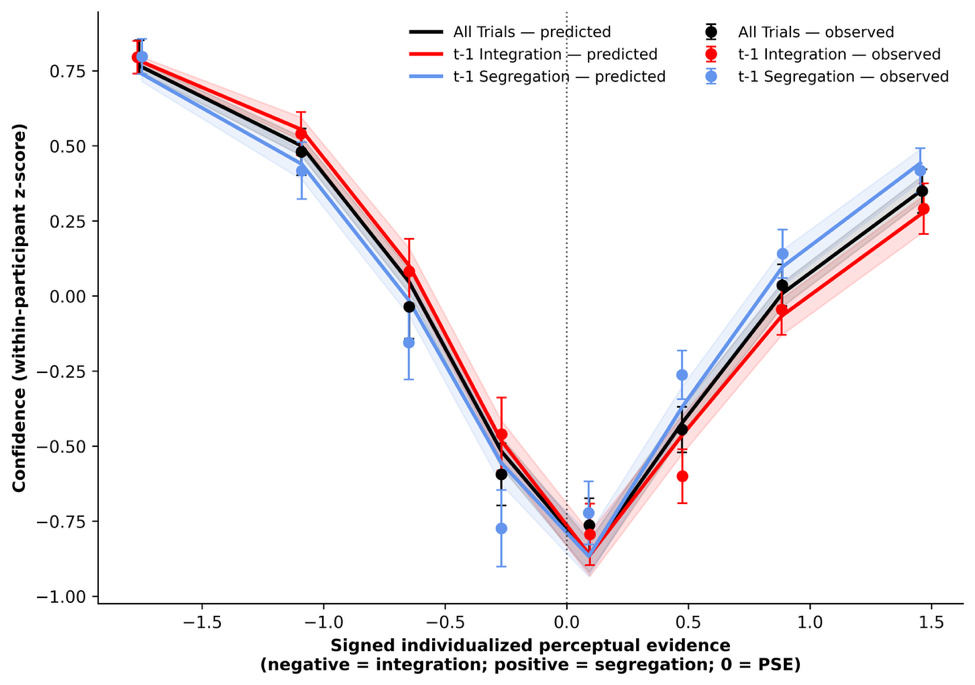
 **Figure S5. Posterior predictive check of confidence across signed perceptual evidence.** Observed and posterior-predicted within-participant standardized confidence are shown as a function of signed individualized perceptual evidence, separately for all trials and for trials preceded by integration and segregation reports. Negative evidence values favor integration, positive values favor segregation, and zero represents the participant-specific perceptual boundary. Solid lines show posterior-predictive medians, shaded regions indicate 95% highest-density intervals, and points with error bars show observed participant-weighted means and 95% confidence intervals. The model reproduced the overall U-shaped relationship between confidence and perceptual evidence, including reduced confidence near the perceptual boundary and increasing confidence with stronger evidence. Local discrepancies remained in several evidence bins, particularly following segregation reports.

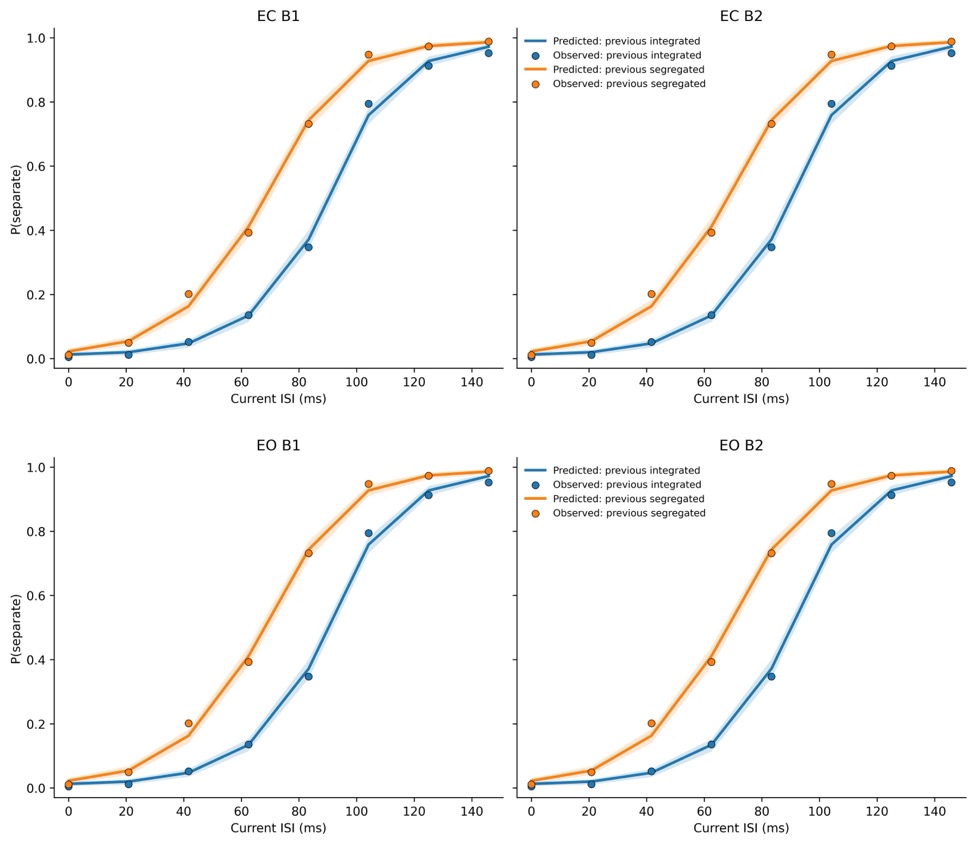
 **Figure S6. Posterior-predictive checks for the IAF-only and IAF + distance models.** Posterior-predicted psychometric functions are shown together with the observed proportion of “separate” responses as a function of current ISI, separately for eyes-closed (EC; top row) and eyes-open (EO; bottom row) resting-state IAF. The left panels show the IAF-only model (B1), in which resting IAF predicts participant-level PSE, whereas the right panels show the extended IAF plus distance model (B2), which additionally includes stimulus frequency–IAF distance. Blue curves and points correspond to trials preceded by an integrated percept, and orange curves and points to trials preceded by a segregated percept. Solid lines show posterior-predicted response probabilities, shaded bands indicate posterior-predictive uncertainty, and circles show the observed group-level response proportions. Both models closely reproduced the observed psychometric functions and the history-dependent shift in temporal-integration thresholds, with highly similar predictions before and after inclusion of the stimulus frequency–IAF distance term.
